# Mechanical Regulation of Microvascular Angiogenesis

**DOI:** 10.1101/2020.01.14.906354

**Authors:** Marissa A. Ruehle, Emily A. Eastburn, Steven A. LaBelle, Laxminarayanan Krishnan, Jeffrey A. Weiss, Joel D. Boerckel, Levi B. Wood, Robert E. Guldberg, Nick J. Willett

**Author notes:** **Address correspondence to:** Nick J. Willett, Emory University School of Medicine, Division of Orthopaedics, 1670 Clairmont Rd, Room 5A125, Decatur, GA 30033.

## Abstract

Neovascularization is a critical early step toward successful tissue regeneration during wound healing. While vasculature has long been recognized as highly mechanosensitive (to fluid shear, pulsatile luminal pressure, etc.), the effects of extracellular matrix strains on angiogenesis are poorly understood. Previously, we found that dynamic matrix compression *in vivo* potently regulated neovascular growth during tissue regeneration; however, whether matrix deformations directly regulate00 angiogenesis remained unknown. Here, we tested the effects of load initiation time, strain magnitude, and mode of compressive deformation (uniform compression vs. compressive indentation that also introduced shear stress) on neovascularization and key angiogenic and mechanotransduction signaling pathways by microvascular fragments *in vitro*. We hypothesized that neovascularization would be enhanced by delayed, moderate compression and inhibited by early, high magnitude compression and by compressive indentation. Consistent with our hypothesis, early, high magnitude loading inhibited vessel growth, while delayed loading enhanced vessel growth. Compressive indentation led to longer, more branched networks than uniform compression – particularly at high strain magnitude. Gene expression was differentially regulated by time of load initiation; genes associated with active angiogenic sprouts were downregulated by early loading but upregulated by delayed loading. Canonical gene targets of the YAP/TAZ mechanotransduction pathway were increased by loading and abrogated by pharmacological YAP inhibition. Together, these data demonstrate that neovascularization is directly responsive to dynamic matrix strain and is particularly sensitive to the timing of load initiation. This work further identifies putative mechanoregulatory angiogenic mechanisms and implicates a critical role for dynamic mechanical cues in vascularized tissue regeneration.

**Statement of Significance:** Mechanical cues influence tissue regeneration, and although vasculature is known to be mechanically sensitive, remarkably little is known about the effects of bulk extracellular matrix deformation on the nascent vessel networks found in healing tissues. Here, we demonstrated that load initiation time, magnitude, and mode all regulate microvascular growth, as well as upstream angiogenic and mechanotransduction signaling pathways. Across all tested magnitudes and modes, microvascular network formation and upstream signaling were powerfully regulated by the timing of load initiation. This work provides a new foundational understanding of how extracellular matrix mechanics regulate angiogenesis and has critical implications for clinical translation of new regenerative medicine therapies and physical rehabilitation strategies designed to enhance revascularization during tissue regeneration.

## Introduction

Vasculature is an abundant and vital component of nearly all tissues, and revascularization is a critical step in the wound healing cascade (1). Angiogenesis, the primary mode of new vessel formation in wound healing, begins as new vessels sprout from adjacent intact blood vessels. New vessel sprouts invade the wounded area and begin to form microvascular networks within just a few days. Concurrent with early angiogenesis, fibroblasts deposit a collagenous provisional matrix, which appears characteristically granular due to its high density of newly formed capillaries. Granulation tissue is then remodeled into mature tissue through a complex series of chemical and physical cues (2), such as the coordinated expression of growth factors and cytokines (3) and dynamically changing ECM properties (4). Granulation tissue also experiences deformation forces; for example, bone experiences compression (5), ligaments and tendons undergo tension (6), venous ulcers are often treated with compression bandages (7), and even cutaneous wounds experience tension during closure (8). Importantly, these tissue-level forces are known to influence the overall progression of healing and the end stage functional outcomes of the repaired tissue. However, remarkably little is known about the effects of ECM deformation experienced by healing tissues on angiogenesis specifically, despite the fact that vasculature has long been recognized as mechanosensitive (9).

Angiogenesis is a highly coordinated process involving multiple phases, including sprout tip cell selection, in which a subset of endothelial cells become migratory; proteolytic ECM invasion; collective cell migration; and cell proliferation and recruitment. Many of these individual processes are known to be responsive to mechanical stimuli. For example, key molecular regulators of tip cell selection (10) have recently been identified as mechanosensitive (11, 12), and expression and secretion of proteases that are involved in angiogenesis are also mechanosensitive (13, 14). Collective cell migration requires the binding and release of ECM by adhesion molecules such as integrins (15), which act as important mediators of ECM-cell mechanotransduction (16). Intracellular mechanotransduction is required to elicit a cellular response, and recently, yes-associated protein (YAP) and its paralog transcriptional coactivator with PDZ-binding motif (TAZ) have been identified in flow-mediated vascular mechanotransduction (17). In other cellular processes, YAP and TAZ are activated by mechanical stimulation, including extracellular matrix forces, and translocate to the nucleus (18), where they function as transcriptional co-activators (19, 20) to upregulate downstream target genes, including *Ctgf* and *Cyr61* (18, 21) – known regulators of angiogenesis (22). YAP and TAZ can also regulate a number of processes crucial to angiogenesis, including VEGF signaling (23), cell migration (21), proliferation, and cell-cell junction formation (24). While many stages of angiogenesis have exhibited mechanosensitivity when investigated in isolation, little is known about how strain from extracellular matrix deformation regulates the concerted processes of angiogenesis during wound healing.

The bone regeneration environment is one of few in which the effects of tissue deformation on neovascularization have been investigated (25–27). Previous work from our lab has shown that compressive deformation of the extracellular matrix (ECM) *in vivo* can have potent effects on neovascular growth – either inhibitory or stimulatory depending on the timing of load initiation. In a critical size segmental bone defect, compressive loading of the defect was applied either immediately following the defect creation surgery, “early”, or four weeks after the surgery, “delayed”. When compression was applied early, concurrent with granulation tissue formation, angiogenesis and subsequent bone tissue formation were significantly inhibited. However, when compressive deformation was delayed and thus applied to newly mineralized, callus-like tissue, blood vessel growth and subsequent healing were enhanced (25), suggesting that ECM deformation regulates angiogenesis in a manner dependent on initiation time.

While the timing of loading has been implicated as an important regulator of ECM deformation effects on angiogenesis, the previously studied *in vivo* bone injury environment is highly complex, and the roles of other deformation parameters such as load frequency, magnitude, and mode (e.g. compression vs. shear) were not well defined. Ambulatory loading typically occurs with a frequency around 1 Hz (28), and strain magnitude is an important regulator of bone formation. Peak bone tissue regeneration occurs at low strains, whereas a fibrous response occurs at high strains; 10% strain is thought to be an approximate transition point between regenerative and non-healing responses (29). Further, while compressive forces are the most widely studied in bone, ambulatory loading includes additional modes of loading such as shear, which has been shown to reduce revascularization of bone (26).

The complexity of the *in vivo* environment hinders comprehensive investigation of the regulatory role played by different loading parameters on the angiogenic healing environment. To exert more precise control over key factors in mechanical loading and matrix deformation and to isolate the vasculature from the complex regenerative environment, we employed an *in vitro* 3D model system of angiogenesis. Microvascular fragments are segments of mature vasculature and composed of multiple cell types, including endothelial cells, smooth muscle cells, mesenchymal stromal cells, and fibroblasts (30, 31). The fragments are typically cultured in collagen-based hydrogels (32), which have similarities to the collagenous granulation tissue characteristic of early stage wound healing. Temporal dynamics of microvascular fragment growth recapitulate the key stages of *in vivo* angiogenesis including sprout tip cell selection, matrix invasion, and neovessel elongation and branching. Additionally, microvascular fragments are known to respond to mechanical cues, including ECM stiffness and tensile deformation (33, 34). Microvascular fragments cultured within collagen-based hydrogels represent a mechanically-sensitive model system to recapitulate the 3D cell-matrix and cell-cell interactions critical to angiogenesis under precisely controlled mechanical loading parameters.

Here, we studied the effects of two load initiation times (early and delayed), three strain magnitudes (5%, 10%, and 30% strain), and two modes of compressive deformation (uniform compression and compressive indentation) on microvascular network growth. We hypothesized that vascularization would be enhanced by delayed, moderate compression and inhibited by early, high magnitude compression. Further, we hypothesized that compressive indentation, which introduced shear stress, would inhibit angiogenesis. Next, we investigated cellular-level changes induced by the timing of load initiation (early vs. delayed): cell viability, cell proliferation, perivascular cell coverage, and gene expression profiles (including specifically targets of the YAP/TAZ mechanotransduction pathway). We found that neovascularization responded directly to dynamic matrix deformation strain magnitude and was particularly sensitive to the timing of load initiation. Gene expression data also showed a divergent response to early vs. delayed loading and implicated a regulatory role for the YAP/TAZ pathway. This work provides a foundational new understanding of how extracellular matrix mechanics regulate angiogenesis, with critical implications for synergizing regenerative therapies and rehabilitation strategies.

## Results

### Non-Loaded Microvascular Fragments Progress Through Distinct Stages of Angiogenesis in vitro

The temporal progression of network formation by microvascular fragments cultured in decorin-supplemented collagen hydrogels was first investigated in the absence of mechanical deformation to establish baseline time points. Under static culture conditions, microvascular fragments formed *in vitro* networks according to a predictable, repeatable time course. At day 0 (i.e. day of harvest), the freshly isolated fragments had characteristically rounded ends, denoted by white arrows in Figure S1. By day 3 in culture, the fragment ends adopted a pointed appearance, indicative of early sprouting and invasion of the extracellular matrix. By day 5, the initial sprouts extended and began to branch. Existing sprouts continued to elongate between days 5-7, and secondary branching began between days 7-10. Sprout and branch initiation processes occurred within the first five days of culture, while days 5-10 consisted primarily of elongation. Thus, days 0-5 were selected as “early loading” to coincide with initiation processes of angiogenesis and days 5-10 were selected as “delayed loading” to coincide with elongation processes (Figure 2 A).

**Figure 1.**
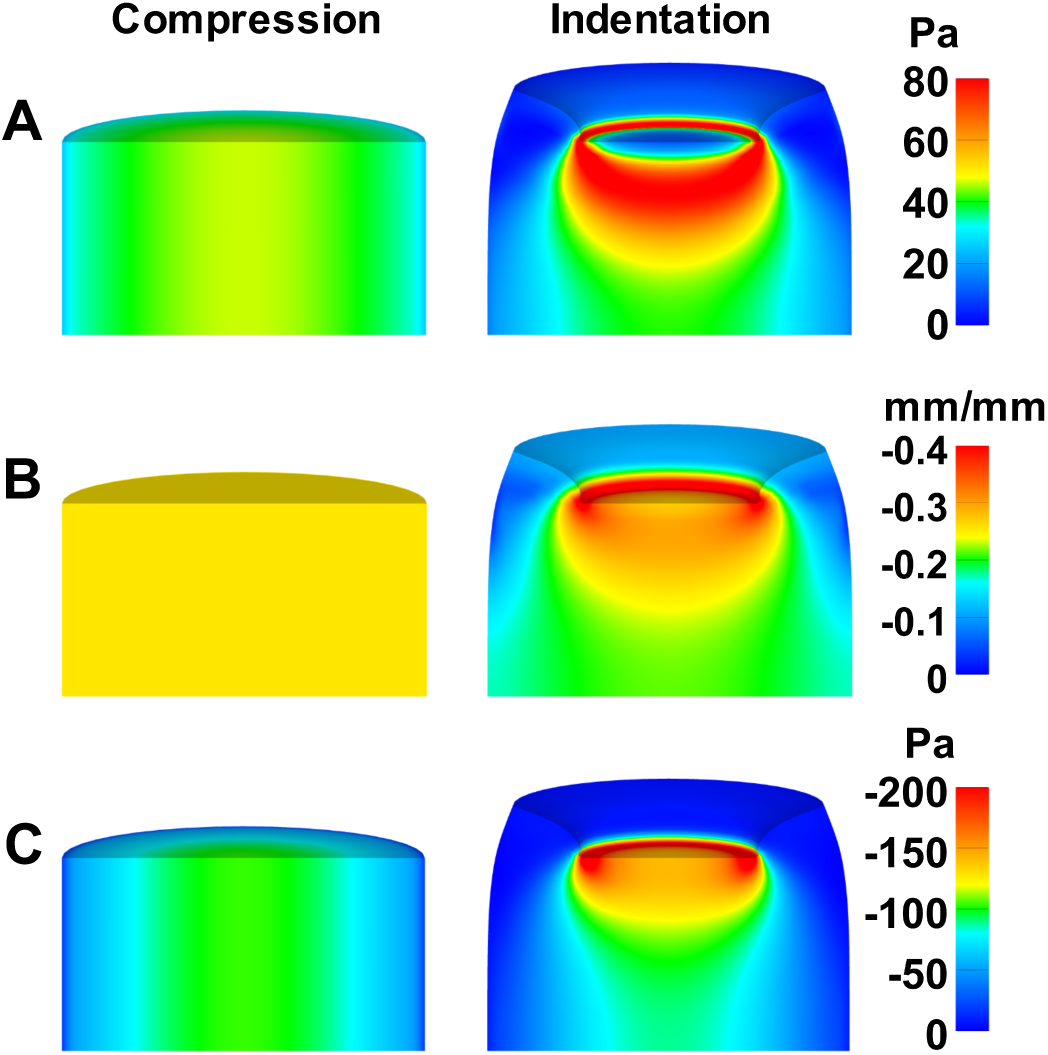
Simulation of decorin-supplemented collagen hydrogels loaded to 30% peak strain based on biphasic constitutive model showing A) maximum shear stress (octahedral; Pa), B) maximum compressive strain (3rd principal strain; mm/mm), and C) maximum compressive stress (3rd principal stress; Pa). Gels loaded in compressive indentation exhibited large stress and strain gradients throughout the gel, whereas gels loaded in uniform compression exhibited virtually no spatial gradients.

**Figure 2.**
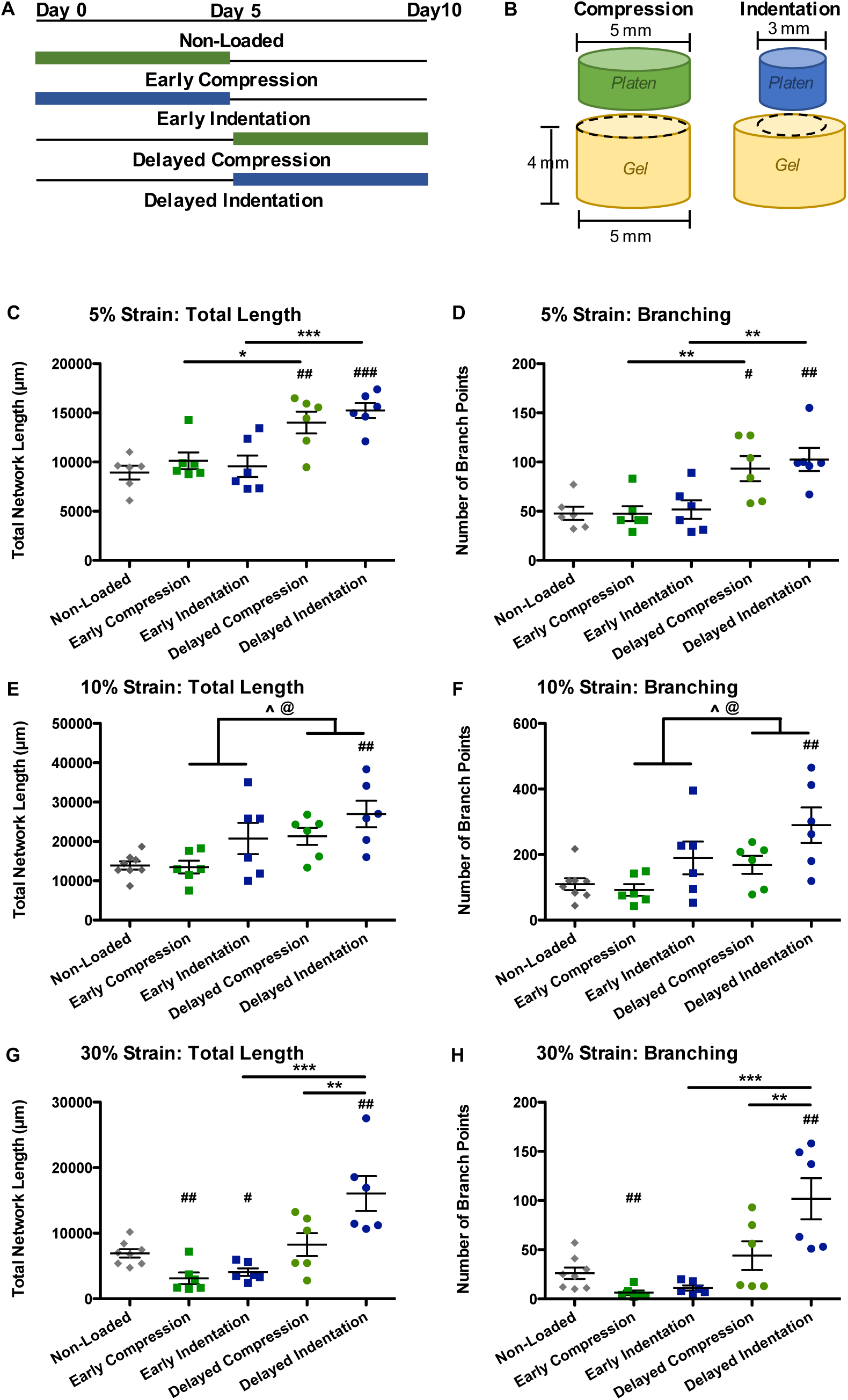
Quantification of MVF network length and branching under strain. A) Microvascular fragments were cultured in decorin-supplemented hydrogels. Gels were loaded continuously for either the first five days of culture (early loading) or the final five days of culture (delayed loading). Gels were loaded at either 5%, 10%, or 30% strain. B) Gels were loaded in compression with platens having a diameter greater than that of the gel or in compressive indentation by platens with a diameter less than that of the gel that also introduced shear stress. C-H) Quantification of microvascular network length and branching based on 3D confocal z-stacks of 200 um depth. * significant difference from NL; 1-way ANOVA, * p<0.05, ** p<0.01, *** p<0.001. # 2-way ANOVA with Bonferroni post hoc test; @ overall effect of loading type; ^ overall effect of time; post hoc # p<0.05, ## p<0.01, ### p<0.001. n=6/group.

### Microvascular Network Formation Exhibits Sensitivity to Magnitude and Mode of Dynamic Loading

In our previous *in vivo* segmental defect study, early ambulatory loading disrupted vascularization within regenerating bone, whereas delayed ambulatory loading enhanced vascular network formation (25); however, the specific parameters of loading that regulate angiogenesis are poorly defined. Informed by bone regeneration literature, we identified 5% strain as pro-regenerative, 10% as transitional, and 30% strain as inhibitory to healing (29). These loads (5%, 10%, and 30% strain) were applied to microvascular fragment-seeded hydrogels with a triangle wave at a frequency of 1 Hz, corresponding to typical gait frequency (28). Gels were loaded in compression with one of two platen configurations: platens having a diameter greater than that of the gel to induce uniform compression (denoted as “compression”), or by platens with a diameter less than that of the gel (denoted as “indentation”; Figure 2 B), to introduce shear along with compression. Computational modeling revealed that the indentation platens created regions of shear stress twice as great as that induced by compression platens (Figure 1 A). Loading was continuously applied either early, days 0-5 of culture, or was delayed until days 5-10 of culture (Figure 2 A). Dynamic loading experiments included the following groups (n=6/group) at 5%, 10%, and 30% strain: early compression, early indentation, delayed compression, delayed indentation, and a non-loaded control.

At 5% strain, delayed loading led to significantly greater total network length (two-way ANOVA, Bonferroni post hoc p<0.05) and number of branches (post hoc p<0.01) compared to early loading (Figure 2 C-D). There was no effect of loading mode at 5% strain (i.e. compression vs. indentation) and no significant interaction effect. Delayed 5% loading, both compression and indentation, increased length (one-way ANOVA, Bonferroni post hoc p<0.01 and p<0.001, respectively) and branching (post hoc p<0.05 and p<0.01, respectively) relative to the non-loaded control. Early 5% loading was not significantly different than the non-loaded control.

At 10% strain, delayed loading again significantly increased total network length (two-way ANOVA, overall effect p<0.05) and number of branches (overall effect p<0.05) relative to early loading (Figure 2 E-F). At 10% strain, indentation also increased length (overall effect p<0.05) and branching (overall effect p<0.05) relative to compression. There was no significant interaction effect. Delayed indentation loading increased total length (one-way ANOVA, Bonferroni post hoc p<0.01) and number of branches (post hoc p<0.01) compared to the non-loaded control. Early 10% loading was no different than the non-loaded control.

At 30% strain, delayed loading significantly increased total network length and number of branches compared to early loading, which was especially pronounced in the indentation groups (two-way ANOVA, Bonferroni post hoc p<0.001; Figure 2 G-H). Delayed indentation increased length (post hoc p<0.01) and branching (post hoc p<0.01) relative to delayed compression. There was no significant interaction effect. Further, delayed indentation significantly increased length (Kruskal-Wallis, Dunn’s post hoc p<0.01) and branching (post hoc p<0.01) over the non-loaded control. At 30% strain, early loading, both compression and indentation, decreased the total network length relative to the non-loaded control (Kruskal Wallis, Dunn’s post hoc p<0.01 and p<0.05, respectively). Early compression also decreased branching as compared to the non-loaded control (post hoc p<0.05). Qualitatively, the early loaded constructs at day 10 primarily exhibited early stage sprouts (Figure 3 A) more comparable to the non-loaded sprouting observed at day 3 (Figure S1).

**Figure 3.**
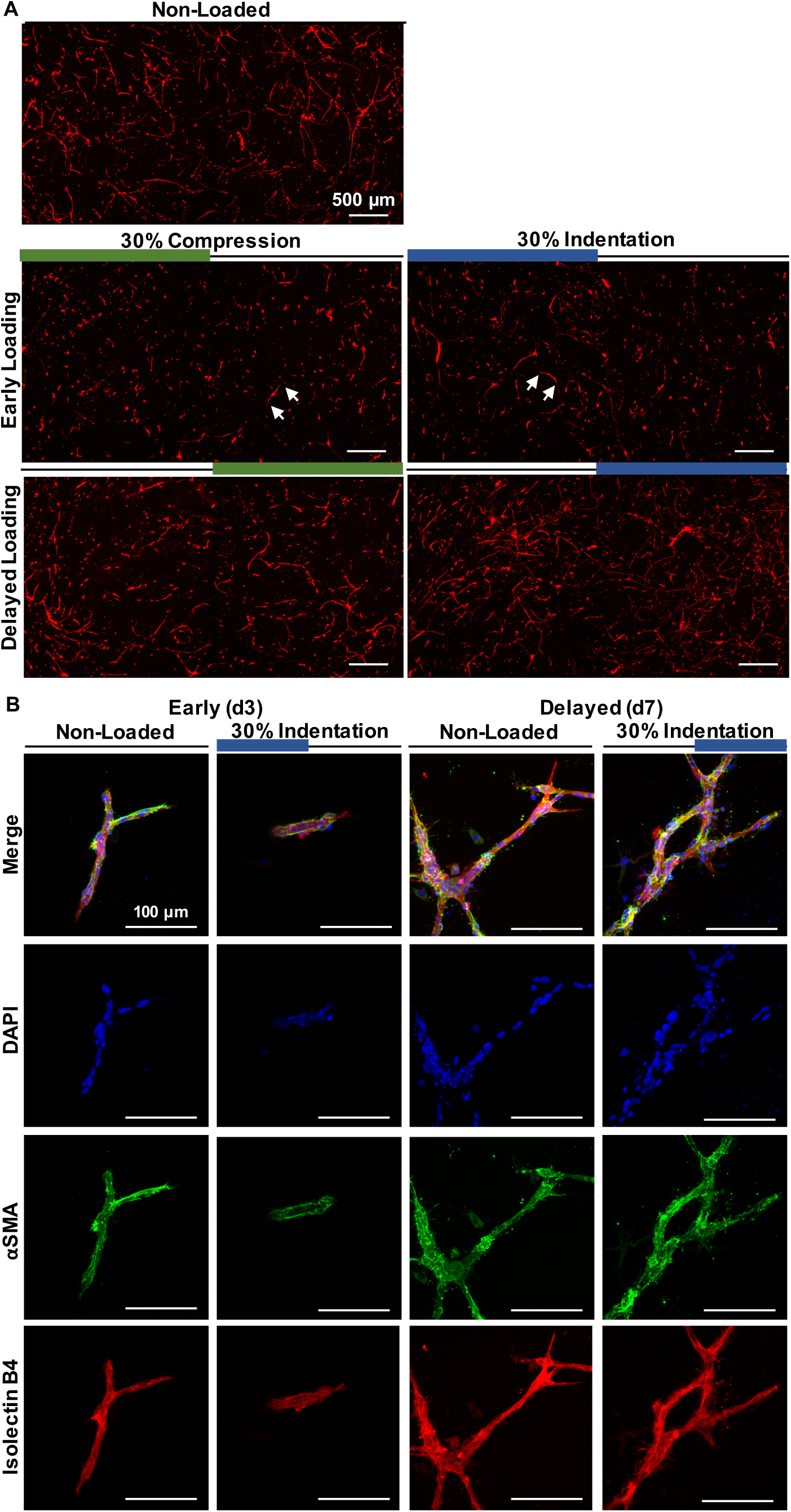
Representative images of non-loaded microvascular networks and networks formed under early vs. delayed dynamic 30% strain at 1 Hz. A) At 30% strain, early compression and early indentation groups exhibit only very early stage sprouts (white arrows), whereas delayed indentation appears qualitatively more densely vascularized than the non-loaded or delayed compression groups. Maximum intensity z-projections (200 um depth) of samples stained with GS-1 lectin at day 10 of culture. Scale bar = 500 µm B) Despite differences in vessel network length and branching in early vs. delayed 30% indentation, microvascular fragments retained perivascular coverage under both early and delayed 30% indentation. Representative maximum intensity z-projections (25 µm depth) of DAPI (blue; nuclei), αSMA (green; perivascular SMCs), and isolectin B4 (red; ECs) stained microvascular fragments at days 3 (early) and 7 (delayed). Scale bar = 100 µm

When all loaded groups’ length and branching were normalized to that of their respective non-loaded experimental controls, early loading exhibited significant strain magnitude-dependence (Figure S2). 30% compression decreased network length and branching relative to both 5% (one-way ANOVA, Bonferroni post hoc p<0.01) and 10% compression (post hoc p<0.05). 30% indentation decreased network length and branching relative to 10% indentation (post hoc p<0.05). There were no statistically significant differences due to strain magnitude among delayed loading conditions. Together, these data demonstrated that delayed loading led to longer, more extensively branched microvascular networks than early loading over a wide range of strain magnitudes.

### Time of Dynamic Loading Initiation Differentially Affects Proliferation but not Viability or Perivascular Cell Attachment

Many cellular-level responses may drive the observed changes in microvascular network morphology, including viability, proliferation, and cell-cell attachments. Based on our results comparing strain magnitude and type of loading, we selected early and delayed 30% strain indentation, the loading parameters that led to the greatest network morphology differences, for subsequent analyses of perivascular cell attachment, cell viability, and proliferation.

As a measure of vascular integrity (35), the spatial relationship between αSMA+ perivascular cells and vessel endothelial cells was assessed at various time points. There were no significant differences in perivascular coverage of endothelial cells due to either early or delayed loading at any time point, and perivascular coverage was approximately 75-80% from day 0 through day 10 (Figure S3 C). Qualitatively, there was relatively less perivascular coverage at the ends of nascent sprouts across all groups.

To assess the effects of dynamic loading on cell viability, a live/dead stain was performed on non-loaded and loaded samples on days 3 (early loading) and 7 (delayed loading) of culture (Figure S3 A). There was no effect of loading on viability, and viability was greater at day 7 of culture than at day 3 for both non-loaded (two-way ANOVA, Bonferroni post hoc p<0.01) and loaded samples (post hoc p<0.01; Figure S3 B). Viability was approximately 75% at day 3 and 90% at day 7.

Proliferation was measured by cellular EdU incorporation also at days 3 and day 7 (Figure 4). There was no effect of early loading on proliferation, and approximately 5-10% of cells were proliferative at day 3. Proliferation was greater at day 7 (two-way ANOVA, overall effect p<0.001), and delayed loading led to increased proliferation compared to non-loaded controls (two-way ANOVA, Bonferroni post hoc p<0.05). Approximately 25% of cells were proliferative at day 7 under delayed loading, while approximately 15% of cells were proliferative at day 7 non-loaded. There was a significant disordinal interaction effect (p<0.05), indicating that early loading and delayed loading have opposite effects on cellular proliferation (Figure QStain B). These data suggest that early and delayed loading differentially affect proliferation, with delayed loading having a stimulatory effect and early loading having a dampening effect. The proliferation results support the observed changes in length and branching; however, the underlying biological mechanisms for these changes remained unclear.

**Figure 4.**
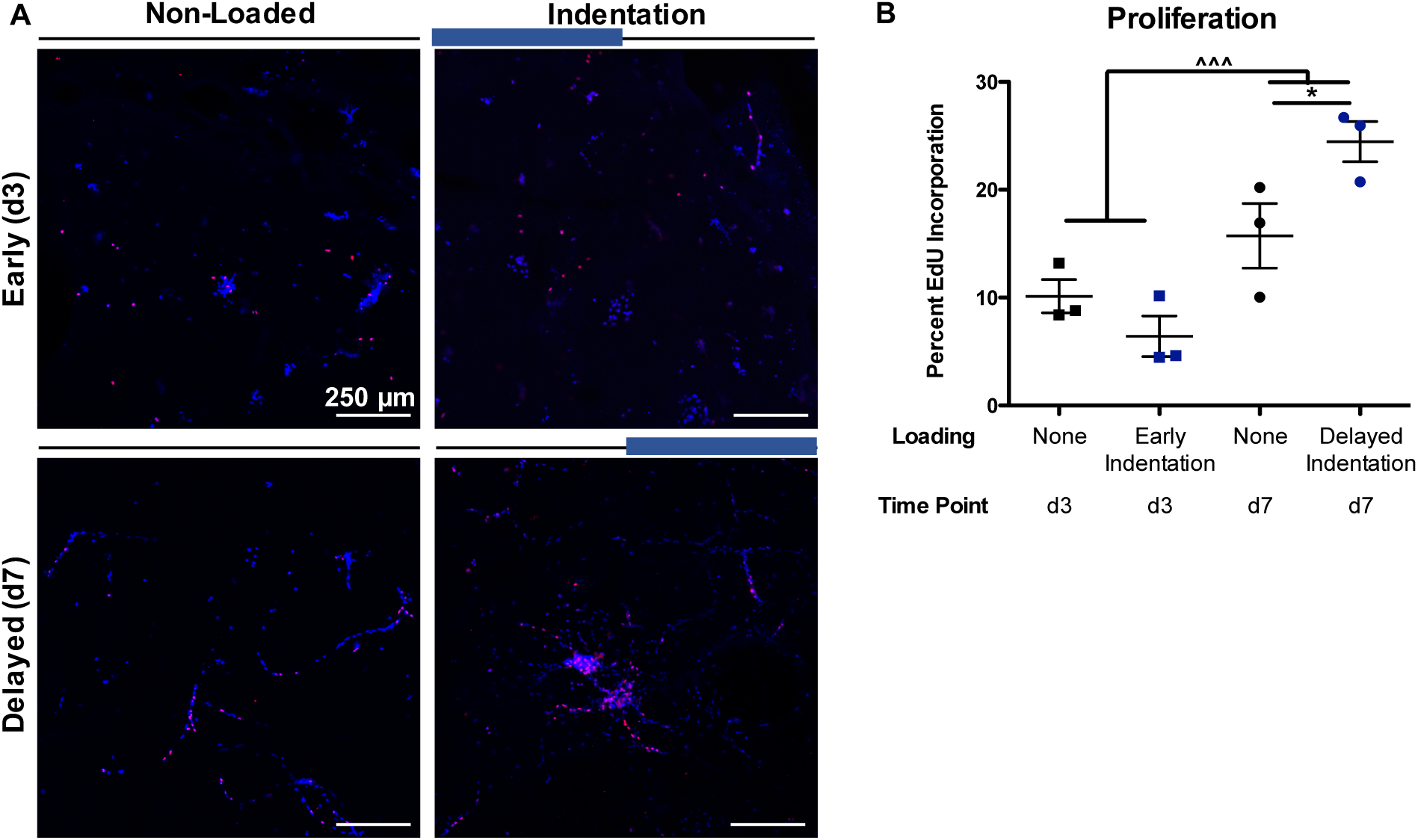
Early vs. delayed 30% indentation loading differentially affected cell proliferation within microvascular networks. Delayed loading increased the number of proliferating cells, and a significant interaction effect suggests that early loading decreased cell proliferation. A) Representative maximum intensity z-projections (25 um depth) of DAPI (blue; all cells) and EdU (red; proliferating cells) stained microvascular fragments at days 3 (early) and 7 (delayed). B) Image-based quantification of proliferation. 2-way ANOVA, ^^^ overall effect of time, p<0.001, post hoc effect of loading ** p<0.01. Significant interaction effect, p<0.05. n=3 gels/group/time point. Scale bar = 250 µm.

### Time of Dynamic Loading Initiation Differentially Regulates Microvascular Fragment Gene Expression

To simultaneously probe the response of multiple key angiogenic processes (e.g. sprout tip cell selection, matrix invasion and deposition, vessel (de)stabilization and growth, adhesion and cell migration, and cell recruitment) to loading, we utilized a high throughput microfluidic RT-PCR gene expression array. Changes in expression of genes related to inflammation, apoptosis, and mechanotransduction were also assessed. Individual genes and corresponding functional sets are shown in Supplemental Table 1; genes and sets were selected based on a literature survey. We focused on early and delayed 30% strain loading – the loading parameters that led to the greatest network morphology differences (Figure S4).

The dimensionality of gene expression data was reduced using partial least squares discriminant analysis (PLSDA) to construct gene expression profiles of non-loaded vs. loaded microvasculature. For both the early and delayed time points, PLSDA generated a latent variable (LV1) that significantly separated non-loaded from loaded samples (1-way ANOVA on score on LV1, post hoc p<0.001; Figure 5 A, C). Latent variables are composed of a weighted average of genes, and each individual gene’s relative contribution can be visualized with a latent variable loading plot. In general, expression levels of many genes across functional groups were downregulated by early loading (negative values in Figure 5 C), while expression levels of many genes across functional groups were upregulated by delayed loading (positive values in Figure 5 D). To analyze the effect of loading on functional gene groups, we performed principal component analysis on each non-overlapping functional gene set (e.g. sprout tip cell selection, matrix invasion and deposition, etc.), then assessed whether the values of principal component 1 significantly differed by group (non-loaded, compression, indentation). We also analyzed the effect of loading on individual genes using one-way ANOVAs.

**Figure 5.**
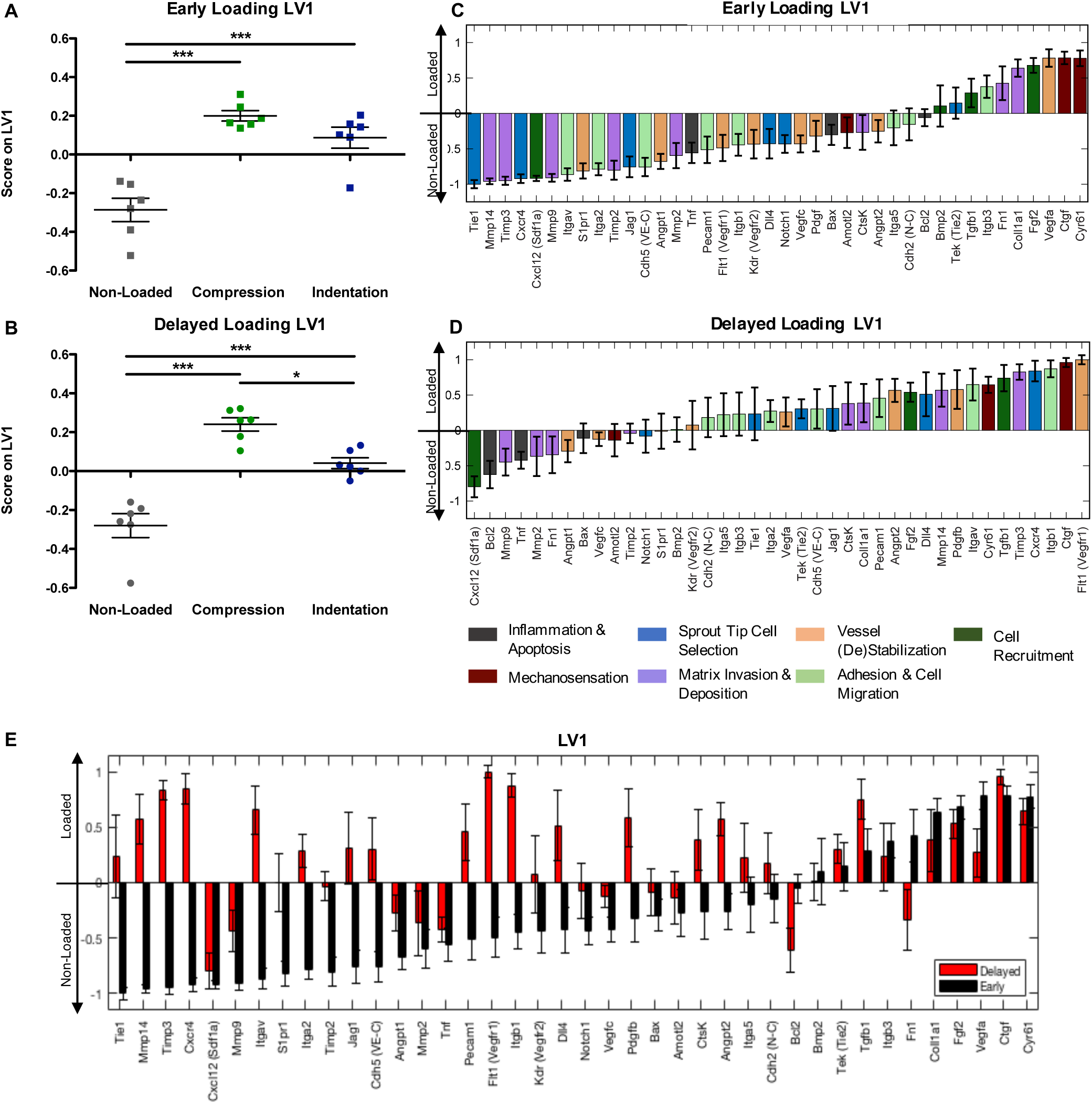
Gene expression array data were analyzed with partial least squares discriminant analysis (PLSDA) to reduce their dimensionality to a latent variable (LV1), which is composed of a weighted average of each individual gene. The gene expression profiles of loaded vs. non-loaded microvasculature, which are represented by LV1, were significantly different for both early (A) and delayed (B) 30% strain loading. Plots of early loading LV1 (C) and delayed loading LV1 (D) demonstrate the weighted average relative contribution of each individual gene to the LV1. In both cases, genes with more positive values are more highly expressed by loaded microvascular networks, and genes with more negative values are more highly expressed by non-loaded microvascular networks. Individual genes are color coded by angiogenic processes (e.g. sprout tip cell selection) they are known to be involved in. E) When interleaved, the loading plots of early loading LV1 vs. delayed loading LV1 reveal differential regulation of a number of genes. Genes strongly downregulated by early loading (negative values in black bars) are often instead upregulated by delayed loading (positive values in red bars), such as *Mmp14*, *Timp3*, and *Cxcr4*. n=6 gels/group. Error bars represent mean±SD of Monte Carlo sub-sampling without replacement.

When considered in aggregate, expression of gene sets known to be involved in sprout tip cell selection (1-way ANOVA on score on principal component 1, post hoc p<0.01) and matrix invasion and deposition (post hoc p<0.01) were significantly downregulated by early loading (Figure S5). Genes associated with matrix invasion (e.g. proteases and protease inhibitors) were also downregulated by early loading whereas matrix deposition genes were upregulated, as evidenced by the loading plot of principal component 1 (Figure S5 D). These data suggest a more quiescent sprout tip cell phenotype may be induced by early loading. Gene sets known to be involved in cell recruitment (post hoc p<0.05) and mechanosensation (overall p<0.05) were upregulated by early loading.

When considered at the individual gene level, *Tie1*, *Mmp14*, *Timp3*, *Cxcr4*, *Cxcl12*, *Mmp9*, and *Itgav* were all significantly downregulated by early loading. *Cyr61*, *Ctgf*, *Vegfa*, and *Fgf2* were all significantly upregulated by early loading (Figure S7). *Tie1*, an orphan receptor that regulates angiogenic sprouting through *Angpt/Tie2* signaling, is expressed by active sprout tip cells and is strongly downregulated in quiescent endothelial cells (36), suggesting that early loading shifts endothelial cells to a more quiescent state. Two other genes associated with tip cells were also downregulated by early loading: *Cxcr4* and *Mmp14*. *Cxcr4* is expressed by activated tip cells (37) and may also play a role in sprout anastomosis (10). The ligand for *Cxcr4*, *Cxcl12* or *Sdf1*, was also downregulated by early loading. *Mmp14* is expressed by tip cells that lead the invasion of surrounding matrix (10, 38). *Timp3* is able to inhibit all MMPs (39), and the fact that it was also downregulated by early loading suggests that the homeostatic balance of MMP-Timp activity may be perturbed by early loading. Although early loading was primarily characterized by a downregulation of gene expression, *Cyr61*, *Ctgf*, and *Vegfa* were strongly upregulated. While *Vegfa* is a necessary component of the sprouting process, it alone is not sufficient to induce sprouting; the balance of other factors, especially angiopoietins 1 and 2, is also a key determinant of whether angiogenesis will occur (40). The increase in *Vegfa* may be a compensatory response of microvascular fragments pushed into a more quiescent state by early loading. The overall increase in cell recruitment genes including *Fgf2* may also reflect a compensatory response. Alternatively, there is evidence that endothelial cells can produce an anti-angiogenic isoform of *Vegfa* (41). The two genes most strongly upregulated in response to early loading were *Ctgf* and *Cyr61*, which are both canonical targets of the YAP mechanotransduction pathway (20).

When considered in aggregate, gene sets known to be involved in cell recruitment (ANOVA on principal component 1, post hoc p<0.001) and mechanosensation (post hoc p<0.01) were upregulated by delayed loading (Figure S6). At the individual gene level, *Flt1*, *Ctgf*, *Itgb1*, *Cxcr4*, *Timp3*, and *Tgfb1* were all significantly upregulated by delayed loading, and *Cxcl12* was significantly downregulated by delayed loading (Figure S8). The five genes most strongly upregulated by delayed loading were *Flt1* or *Vegfr1*, *Ctgf*, *Itgb1*, *Cxcr4*, and *Timp3*. By increasing expression of *Vegfr1*, delayed loading may increase the sensitivity of microvascular fragments to pro-angiogenic Vegf signaling. This is in contrast with early loading, in which *Vegfa* upregulation was not accompanied by *Vegfr1* or *Vegfr2* upregulation. Delayed loading led to strong upregulation of both *Cxcr4* and *Timp3*, which is also in contrast with early loading. Although *Cxcr4* was differentially affected by early vs. delayed loading, its ligand *Cxcl12* or *Sdf1a* was strongly downregulated by both loading scenarios. Although *Cxcl12* can function as a potent cell recruitment molecule, it may play a different role in our system. *Tgfb1* is also considered a potent cell recruitment signal and was upregulated by delayed loading. *Itgb1* is an adhesion molecule essential for angiogenesis (42).

Delayed compression and delayed indentation were also significantly separated along LV1 (1-way ANOVA, post hoc p<0.05; Figure 5 B). Of the 43 genes tested, only *Itga2* expression was significantly higher in indentation relative to compression (one-way ANOVA, Bonferroni post hoc p<0.05) and to the non-loaded control (post hoc p<0.01), despite the large morphological differences observed at day 10. Integrin subunits a2 and b1 can form a complex that can bind both collagen (43) and decorin (44) – the two components of our gel matrices.

A number of genes were differentially regulated by early vs. delayed loading (Figure 5 E); the strongest downregulated contributors to early loading LV1 were instead upregulated by delayed loading, and the strongest upregulated contributors to delayed loading LV1 were instead downregulated by early loading (e.g. *Tie1*, *Mmp14*, *Timp3*, *Flt1* or *Vegfr1*, *Itgb1*, *Cxcr4*). There were two strongly upregulated contributors to both early LV1 and delayed LV1: *Ctgf* and *Cyr61*, which are canonical targets of the YAP mechanotranduction signaling pathway (20).

### YAP is Involved in Microvascular Response to Delayed Loading

The genes with the largest induction by mechanical loading, regardless of loading mode, were *Ctgf* and *Cyr61*. Early loading led to a nearly three-fold increase in *Cyr61* expression for both compression and indentation, and delayed loading led to a 3.4-fold and 5-fold increase in *Ctgf* expression for compression and indentation, respectively. Notably, *Ctgf* and *Cyr61* are both canonical target genes of the mechanosensitive transcriptional co-activators YAP and TAZ (18). To determine whether the mechanoactivation of *Ctgf* and *Cyr61* was indeed YAP/TAZ-dependent, we employed a pharmacological inhibitor of YAP, verteporfin (VP; 5 µM). VP has been shown to inhibit YAP/TAZ activity by disrupting YAP’s formation of a transcriptional complex with TEAD (45) and by sequestering YAP in the cytoplasm, thus preventing nuclear translocation (46). Thus, we hypothesized that the addition of VP would abrogate the increased expression of target genes *Cyr61* and *Ctgf* due to loading. Since the gene expression profiles of compression and indentation were nearly identical in our initial array, only indentation was studied with VP.

At the early time point, loading increased expression of *Ctgf* (overall effect p<0.01; Figure 6), but there was no statistically significant effect of VP on early gene expression of either *Ctgf* or *Cyr61*. At the delayed time point, delayed loading without VP increased expression of both *Ctgf* (two-way ANOVA, Bonferroni post hoc p<0.001) and *Cyr61* (post hoc p<0.001) relative to the non-loaded control. VP significantly abrogated the increased expression of *Ctgf* (post hoc p<0.05) and *Cyr61* (post hoc p<0.001) induced by loading, suggesting that YAP mediates the mechanotransductive gene induction by delayed loading. Interestingly, a similar abrogation of *Ctgf* and *Cyr61* expression by VP was not observed in early loaded samples despite their induction in the initial gene expression array.

**Figure 6.**
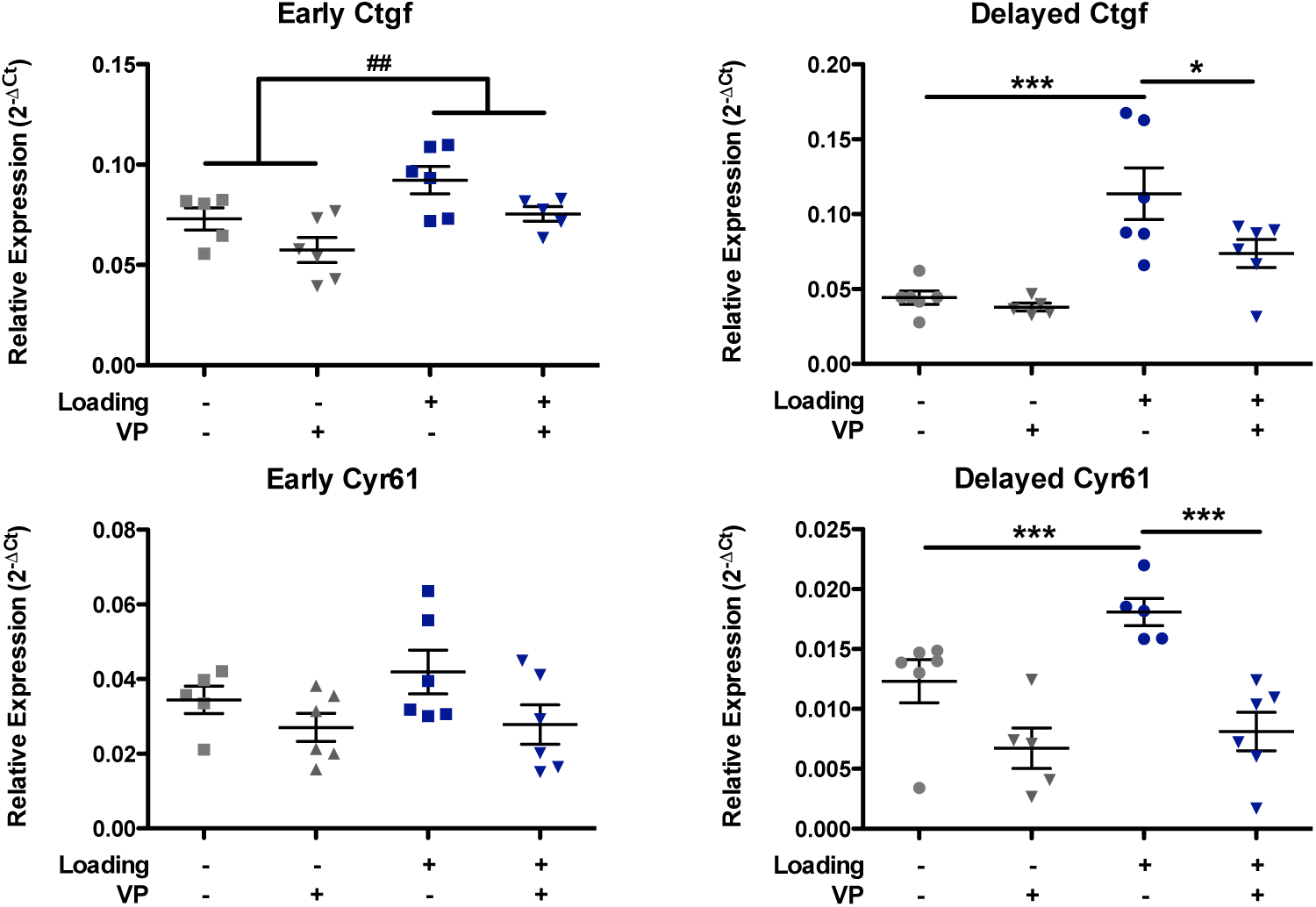
Expression of YAP/TAZ target genes *Ctgf* and *Cyr61* relative to mean expression of housekeeping genes. Early 30% indentation loading induced significant upregulation of *Ctgf*, and delayed 30% indentation loading induced significant upregulation of both *Ctgf* and *Cyr61*. YAP inhibitor, verteporfin (VP, 5 µM) abrogated the delayed loading-induced upregulation of both target genes. 2-way ANOVA, overall effect of loading ## p<0.01; Bonferroni post hoc *p<0.05, **p<0.01, ***p<0.001. Delayed *Ctgf* demonstrated significant interaction effect. n=5-6/group.

## Discussion

In this set of studies, we demonstrated that dynamic extracellular matrix deformation profoundly influences microvascular network formation and that the initiation time, mode, and magnitude of compressive strain are all critical regulatory parameters. Across all conditions tested, delayed compressive loading (initiated at day 5 following initial sprout and branch initiation) enhanced vessel network formation compared to early loading (initiated at day 0). These morphological differences (i.e. longer, more extensively branched networks due to delayed vs. early loading) were mirrored by increased cell proliferation in response to delayed loading and divergent regulation of genes associated with active angiogenic sprouts, where many of the same genes were downregulated by early loading but upregulated by delayed loading. Together, these data implicate the timing of load initiation as a critical determinant of vascular network formation and suggest that the early stages of angiogenic sprouting are exquisitely sensitive to the bulk ECM deformation that would be experienced by healing tissues.

Load magnitude is often implicated as the parameter of mechnoregulation that primarily dictates regenerative responses. However, our previous results have shown that the timing of load application, even loads of similar magnitudes, has a profound effect. Consistent with our hypothesis and with prior *in vivo* results (25, 47), delayed loading produced greater network length and number of branches at all strain magnitudes tested. At low strains (5% and 10%), early loading did not affect vessel network formation; however, at a much higher strain (30%), early loading significantly decreased network length and branching. These data suggest that low magnitude strain may be permissive to early stage angiogenesis, whereas higher magnitude strains can disrupt this process. Inhibitory effects of delayed loading were not observed, even at 30% strain, suggesting that once a critical stage of vascular network maturity is reached, even high magnitude loading is permissive to vessel growth. Day 5 was chosen as the inflection point of early vs. delayed loading in the present studies to distinguish between the distinct processes of sprout and branch initiation vs. elongation, suggesting that the initial formation of sprouts/branches may be this critical stage.

In addition to load magnitude, previous work has also investigated the effects of different magnitudes of loading, and shear stress has been shown to be detrimental to neovascularization of regenerating bone tissue (26). Thus, we hypothesized here that shear stress may directly inhibit angiogenesis. Contrary to our hypothesis, compressive indentation that also introduced shear stress increased vessel length and branching overall compared to compression alone at strains above 5%. Previous work that demonstrated a reduction in bone neovascularization due to shear focused on the effects of translational shear, i.e. perpendicular to the axis of compression (26), whereas in the current study we utilized indentation platens that were designed to introduce a zone of interfacial shear. Although there were no differences observed in the microvascular networks at the interface of contact between the platen and gel and there was no difference in the spatial distribution of vessels, the indentation platens did indeed introduce shear stress of up to twice the magnitude of that introduced by the compression platens in the center of the gels (Figure 1 A). However, the indentation platens also profoundly changed the local strain and stress gradients (Figure 1 B-C) producing high strain gradients throughout the gels, which are known to also influence tissue-level remodeling. A previous study similarly found that new bone tissue formation can localize specifically at sites of high strain gradients (48). In contrast, there was virtually no strain gradient within the uniformly compressed gels. The effects of strain gradients may also explain the differing effects of strain magnitude in uniform compression vs. compressive indentation. Delayed compressive indentation increased vascular network length and branching at all three tested strain magnitudes, including the high 30% strain. In contrast, delayed uniform compression exerted a significant stimulatory effect at 5% strain, a positive significant overall effect but not statistically significant post-hoc difference at 10% strain, and no effect at 30% strain. The trend observed in uniform compression was more consistent with our hypothesis and the bone regeneration literature focused on strain magnitude; lower strains tend to promote bone regeneration, while 30% compressive strain is sufficiently high that it is not often found under physiologic conditions and disrupts bone healing (29, 49, 50). Our work suggests a critical role for strain gradients in regulating angiogenic processes and motivates further investigation of the effect of strain gradients on enhancing vascularization. Future computational work may more precisely define these effects.

Although the delayed compression and delayed indentation environments produced differences in terms of vascular network length and branching, gene expression from the 43 genes tested in our array was similar between the conditions, with only one (*Itga2*) showing differential expression between the conditions. The lack of differences in gene expression profiles between these groups may be due to the sampling of a single time point after 24 hours of loading; gene expression, and subsequent morphologies, may diverge later between loading conditions. Future work may also include an unbiased analysis of additional genes using techniques such as RNA-seq. Alternatively, the similar gene expression profiles may suggest that later (after 5 days of loading rather than 24 hours) differences may be primarily mediated by parameters that were not assessed in this system, such as convective fluid flow patterns, which are difficult to decouple from *in vitro* loading systems. However, if all effects observed in these experiments were due to an increase in fluid flow and thus nutrient transport, early 30% loading would likely have also had a beneficial effect rather than the inhibitory effect we observed.

Early vs. delayed gene expression results mirrored the morphological results, with early loading resulting in decreased vascular network formation and decreased overall gene expression and delayed loading instead resulting in enhanced vascular network formation and increased gene expression. PCA analysis revealed that expression of the gene set known to be involved in sprout tip cell selection was significantly decreased due to early loading. This suggests that early stage sprout tip cell selection, which is regulated by a balance of Notch interaction with pro-angiogenic *Jag1* and anti-angiogenic *Dll4*, is perturbed by loading. The Notch-Jagged signaling axis has recently been shown to be mechanically sensitive, including the downregulation of pro-angiogenic *Jag1* in response to mechanical stretch (12). In our data, we observed a downregulation of not only *Jag1* but also an overall downregulation of the sprout tip cell selection gene set, including anti-angiogenic *Dll4*. We also observed an overall downregulation of pro-migratory protease expression but an increase in matrix deposition genes. Taken together with the strong downregulation of *Tie1*, characteristic of quiescent endothelial cells (36), our data suggest the induction of a more complex quiescent, non-sprouting phenotype in response to early loading. In contrast, when loading was delayed until after initial sprout tip cell selection has already occurred, we saw an overall upregulation of gene expression – including many of the same genes that were instead downregulated by early loading: *Tie1*, *Mmp14*, *Timp3*, *Flt1* or *Vegfr1*, *Itgb1*, *Cxcr4*. Of these genes, *Tie1*, *Mmp14*, *Flt1* or *Vegfr1*, and *Cxcr4* in particular are known to be expressed by actively migrating sprout tip cells (36–38). Together, these divergent effects of early vs. delayed loading suggest that tip cell activity may be the mechanism through which loading affects neovascularization; early loading depresses sprout tip cell selection signaling, while delayed loading increases expression of genes associated with active tip cells and cell proliferation.

In our mechanically stimulated system, expression of YAP/TAZ target genes *Cyr61* and *Ctgf* was upregulated by both early and delayed loading. Interestingly, although we also observed significant upregulation of *Cyr61* and *Ctgf* in the initial gene expression array, we did not observe the same significant upregulation in the VP experiment. These samples contained DMSO to appropriately control for the delivery of VP within DMSO, and DMSO may have altered the baseline response of microvascular fragments to early loading. The regulatory role played by YAP/TAZ may be determined by other factors concurrently affected by early vs. delayed mechanical loading, and the role of YAP/TAZ in regulating divergent responses requires future work. DMSO has been previously reported to have anti-angiogenic effects (51), which also prevented functional analysis of microvascular network formation in the presence of VP; the authors acknowledge this as a limitation of the present studies. *Itgb1*, an essential adhesion molecule for angiogenesis (42) known to be involved in YAP/TAZ signaling (21), was upregulated by delayed loading. The one molecule that was differentially expressed by delayed compression vs. indentation was also an integrin, *Itga2*. The a2 and b1 integrin subunits form a complex that can bind both collagen (43) and decorin (44), the two components of our gel matrices. In our system, integrins a2b1 may couple cells to the mechanically dynamic matrix and thus act as an element of the mechanotrasduction pathway.

YAP and TAZ have recently emerged as mechanotransducers to regulate critical processes including organ size, tumor growth, and stem cell fate (52). In vasculature, they have recently been implicated in transducing the atherogenic effects of oscillatory shear stress (17) – one of the most well-studied examples of vascular mechanosensitivity (9). Additionally, YAP and TAZ are known to promote sprouting angiogenesis, even in systems that are not directly mechanically stimulated (24), through molecular regulators of sprout tip cell selection (53). In our studies, we observed a strong upregulation of YAP/TAZ target genes *Cyr61* and *Ctgf* due to loading; this response was significantly abrogated by YAP/TAZ inhibitor VP only under the delayed loading condition. These results suggest that delayed loading activates YAP signaling (perhaps through Itgb1), which leads to an increased number of active sprout tip cells (53) and thus upregulation of genes associated with active tip cells, to enhance vascularization.

## Conclusions

Here, we have demonstrated that the magnitude, mode, and initiation time of ECM loading are all critical regulators of angiogenesis. Across all tested magnitudes and modes, delayed loading enhanced vessel network formation relative to early loading. Morphological differences were mirrored by increased cell proliferation in response to delayed loading and divergent regulation (downregulated by early loading and upregulated by delayed loading) of genes associated with active angiogenic sprouts. Together, these data implicate time of load initiation as a critical determinant of vascular network morphology and suggest that the early process of angiogenic sprouting is exquisitely sensitive to bulk loading of their ECM. By providing increased foundational understanding of the time-dependent mechanical regulation angiogenesis, this work begins to enable mechanical loading to be leveraged as a therapeutic component of future tissue engineering and physical rehabilitation approaches.

## Methods

### Microvascular Fragment Isolation and Culture

Microvascular fragments were isolated as previously described (32). Briefly, epididymal fat pads of retired breeder Lewis rats were harvested, minced, and digested in a collagenase solution. Microvascular fragments were obtained through selective filtration to retain multicellular structures between 20-200 µm. The fragments were suspended at a density of 20,000 fragments/mL in 3% collagen gels supplemented with 50 µg/mL decorin (DCN) to improve construct dimensional stability (54). Gels were formed by 15-20 minutes of incubation at 37 °C in custom polycarbonate molds to create gels with a diameter of 5 mm and a height 4 mm. Microvascular fragment-containing gels were cultured in serum-free media supplemented with 10 ng/mL recombinant human vascular endothelial growth factor (rhVEGF; R&D Systems, Minneapolis, MN) (55). Media was changed on days 3, 5, and 7 of culture, and gels were fixed with 4% paraformaldehyde on day 10.

### Dynamic Loading

Microvascular fragment-containing gels were loaded using an Electroforce 5500 with a multi-specimen compression chamber containing a 24-well plate loading assembly (TA Instruments; New Castle, DE). Loading was applied in a triangle wave with amplitudes corresponding to 5%, 10%, or 30% strain (0.2 mm, 0.4 mm, and 1.2 mm, respectively) at a frequency of 1 Hz. Gels were loaded in compression using polyetheretherketone (PEEK) platens with a diameter greater than that of the gel (1 cm) or in compression with a shear interface zone (indentation) using PEEK platens with a 3 mm gel-contacting diameter. Loading was applied continuously, breaking only for media changes, for either the first five days of culture, early loading, or the final five days of culture, delayed loading. To ensure gels remained centered within the well, gels sat within the inner diameter of 1 mm thick 3D printed poly(lactic-co-glycolic acid) rings during loading.

To study the effects of loading mode and time of initiation, dynamic loading at 5%, 10%, and 30% strain experiments included the following groups (n=6/group): early compression, early indentation, delayed compression, delayed indentation, and a non-loaded control.

### Computational Simulation of Dynamic Loading

Stress relaxation data of decorin-supplemented collagen hydrogels from prior publication were fit to a biphasic constitutive model with a neo-Hookean ellipsoidal fiber distribution solid phase (54). The solid volume fraction was approximated based on the effective specific volume of collagen at 3 mg/mL (56, 57). Permeability and fibril modulus were determined using the parameter optimization module of FEBio during a simulation of stress relaxation (58). A 90 degree wedge geometry was meshed with radial biasing away from the center and vertical biasing away from the contact surface to better accommodate high strains. The geometry was divided into 10 circumferential divisions providing 9 ⁰ resolution for each element. Symmetry boundary conditions were enforced by fixing nodes along the x axis in the y direction and nodes along the y direction in the x direction, restricting deformation to lateral directions. Free draining surfaces were modeled by prescribing zero fluid pressure at the radial edge of the wedge and on the exposed portion of the top surface for the case of compressive indentation. Nodes along the bottom of the gel were fixed in the vertical direction. Gels were deformed using rigid body contact to peak strains of 5, 10, or 30% strain. To achieve convergence in the presence of high strain rates dynamic loading was ramped with displacement prescribed at 2/3 of peak strain at 0.2 Hz for 25 cycles, 9/10 of peak strain at 0.2 Hz for 25 cycles, and finally peak strain at 1 Hz until less than a 0.1% change in 3^rd^ principal stress and shear stress was achieved between cycles for all models. The volume average for stress and shear were determined at full depression during the last cycle by weighting the elemental values by the current volume of each element.

### Staining, Imaging, and Image-Based Analyses

To assess network morphology at day 10, fixed gels were stained with rhodamine-labeled Griffonia simplicifolia (GS-1) lectin (Vector Labs, Burlingame, CA) at a concentration of 5 µg/mL in phosphate buffered saline (PBS) overnight at 4 °C (n=6/group). Gels were imaged using a Zeiss 700 confocal microscope with a 5X objective. The entire diameter of each gel was imaged to a depth of 200 µm. Confocal z-stacks were median filtered, deconvolved, and thresholded using Amira for Life Sciences (ThermoFisher Scientific, Waltham, MA). Islands smaller than 30 voxels (e.g. single cells, debris, or noise) were also removed using Amira. Thresholded images were exported for skeletonization and quantification of length and branching using the 4-D open snake method (59) of the Farsight Toolkit (60, 61).

Viability and proliferation of non-loaded vs. loaded microvascular fragments were measured at day 3 for early loading and at day 7 for delayed loading. Viability was determined using a live/dead assay kit performed according to manufacturer’s instructions (ThermoFisher). Gels were imaged at 10X, and three randomly selected fields were imaged per gel to a depth of 25 µm. Maximum intensity z-projections were created to quantify viability using Fiji’s Analyze Particles feature. Percent viability was calculated as the pixel area of live cells (green channel) over the pixel area of all cells (green+red channels). Proliferation was assessed using a Click-iT EdU Alexa Fluor 594 Imaging Kit (ThermoFisher; n=3/group/time point). EdU was added to media at a concentration of 10 µM at days 2 and 6 of culture and incubated for 24 hours before gels were fixed at days 3 and 7. Gels were imaged at 10X to a depth of 25 µm, and four images were taken to capture the diameter of each gel. Proliferation was quantified as the number of EdU+ nuclei over the total number of nuclei.

To assess the degree of perivascular coverage of endothelial cells by smooth muscle cells over time, non-loaded gels were fixed at days 0, 3, 5, 7, and 10 (n=3/time point), early loaded samples were analyzed at days 3, 5, and 10 (n=3/time point), and delayed loaded samples were analyzed at days 7 and 10 (n=3/time point). Following fixation, gels were stained with Alexa Fluor 488-conjugated anti-alpha smooth muscle actin (αSMA) antibody (ab184675, Abcam, Cambridge, UK) at a 1:100 dilution, DyLight 649-conjugated GS-1 isolectin B4 (DL-1208, Vector Labs) at 5 µg/mL, and DAPI (ThermoFisher) at a 1:1000 dilution. Gels were imaged at 40X to a depth of 25 µm, which is the approximate depth of a single vessel. A region of interest (ROI) was drawn around vascular structures, and signal overlap between αSMA and isolectin B4 were quantified for the ROI using Manders coefficients as determined by the Fiji plugin coloc2 (62).

### Gene Expression Analyses

To assess gene expression changes due loading, RNA was harvested from non-loaded and loaded constructs 24 hours after load initiation (i.e. after 24 hours of culture total for early loaded samples and their non-loaded controls and after 6 days of culture total for delayed loaded samples and their non-loaded controls). RNA was collected from non-loaded and loaded gels at both early and delayed time points (n=5-6/group/time point). RNA was extracted using Qiagen MinElute kits, and cDNA was made using Qiagen RT^2^ First Strand kits (Qiagen, Hilden, Germany). RNA concentration as determined by NanoDrop spectrophotometer (ThermoFisher) was used to ensure cDNA concentrations were equivalent. Taqman probes were used to assess gene expression of 43 genes known to be involved in various stages of angiogenesis and five housekeeping genes (Table 1). Gene expression was quantified using a Biomark real-time PCR integrated fluidic circuit array (Fluidigm, South San Francisco, CA). Rat universal cDNA (Gene Scientific, Rockville, MD) and ultrapure water (ThermoFisher) were used as positive and negative controls, respectively. Data were normalized on a per sample basis to the mean of three housekeeping genes that did not have significantly different levels of expression across groups (Gapdh, Ubc, Hrpt1) using the ΔCt method.

### Multivariate Analysis of Gene Expression Data

Partial least squares discriminant analysis (PLSDA) was performed in MATLAB (Mathworks, Natick, MA) using Cleiton Nunes’s partial least squares algorithm (MathWorks File Exchange). To avoid biasing results with the absolute magnitude of different genes’ expression levels, data were z-scored prior to being analyzed with PLSDA. Orthogonal rotations were applied to the z-scores to maximally separate groups (non-loaded, compressive loading, and indentation loading) based on latent variables 1 and 2 (LV1 and LV2) created by the PLS algorithm. LV loading plots show the mean and standard deviation of each gene’s relative contribution to the latent variable; mean and standard deviation were calculated using Monte Carlo sub-sampling that iteratively excluded a randomly chosen 15% subset of the data 1000 times (63).

Principal component analysis (PCA) was performed on non-overlapping gene sets known to be involved in sprout tip cell selection, matrix invasion and deposition, vessel (de)stabilization and growth, adhesion and cell migration, cell recruitment, inflammation and apoptosis, and mechanotransduction using the MATLAB pca command. Loading plots for principal component 1, which explains the greatest amount of variation in the data, represent the relative contributions of each gene within the set.

### YAP Inhibition

The YAP inhibitor verteporfin (VP; MilliporeSigma, Burlington, MA) was dissolved in DMSO and added to serum-free media at a concentration 5 µM. Non-VP controls received an equal volume of DMSO. VP and DMSO were added to microvascular fragment-containing gels 30 minutes prior to initiation of loading to allow diffusion throughout the gel. After 24 hours of loading, RNA was collected from early and delayed samples as above. The groups for the YAP inhibition study were indentation + VP, indentation + DMSO carrier only, non-loaded + VP, and non-loaded + DMSO at both early and delayed time points (n=5-6/group). All culture of samples containing light-sensitive VP (45) was conducted in the dark.

### Statistical Analysis

For day 10 microvascular network morphology data, a two-way ANOVA was used to compare early vs. delayed loading and compression vs. indentation loading. A one-way ANOVA was used to compare early loading to the non-loaded control and delayed loading to the non-loaded control; a Kruskal-Wallis test with Dunn’s post hoc test was used in cases where the variances significantly differed among groups. To directly compare the effects of different load magnitudes, loaded groups were normalized to their respective non-loaded controls and analyzed within time point and loading mode (e.g. early indentation compared at 5%, 10%, and 30% strain) using a one-way ANOVA. Viability, proliferation, and perivascular coverage data were analyzed by two-way ANOVA. Gene array data were studied in aggregate using PLSDA as detailed above, and statistical significance was determined using a one-way ANOVA on each group’s mean score on LV1. The expression levels of individual genes were compared within time point using a one-way ANOVA. The effect of loading on entire gene sets was assessed with PCA as detailed above, and statistical significance was determined using a one-way ANOVA on each group’s mean score on principal component 1. The effect of verteporfin on gene expression of non-loaded vs. loaded constructs was analyzed with a two-way ANOVA. Bonferroni’s post hoc test followed all ANOVAs. All statistical analyses were performed in GraphPad Prism 5 with α=0.05. Data are presented as mean ± standard error of the mean.

## Acknowledgements

This work was supported by funding from the National Institutes of Health (NIH) R01 AR069297. This material is the result of work supported with resources and the use of facilities at the Atlanta VA Medical Center; the contents do not represent the views of the U.S. Department of Veterans Affairs or the United States Government. The authors wish to acknowledge the core facilities at the Parker H. Petit Institute for Bioengineering and Bioscience at the Georgia Institute of Technology for the use of their shared equipment, services, and expertise. We also thank Brett Klosterhoff for assistance with 3D printing and machining.

