## Supplemental Figures for "Mechanical Regulation of Microvascular Angiogenesis"

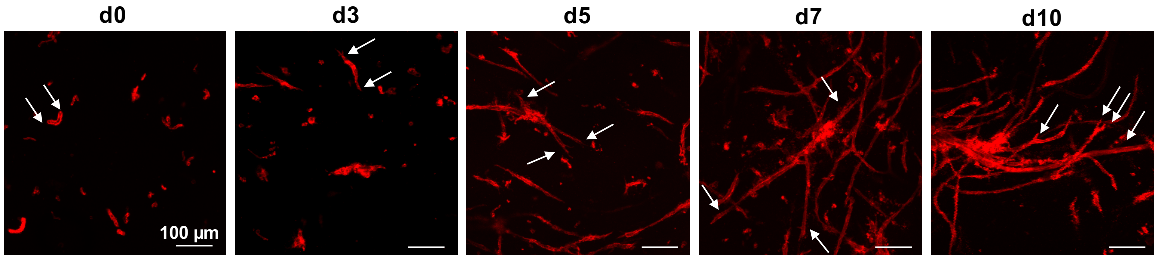


**Figure S1.** Representative images of *in vitro* microvascular network formation over time. White arrows denote ends of freshly isolated fragments (d0) and resulting sprouts (d3, d5, d7, d10). Maximum intensity z-projections (200 µm depth) of samples stained with GS-1 lectin.

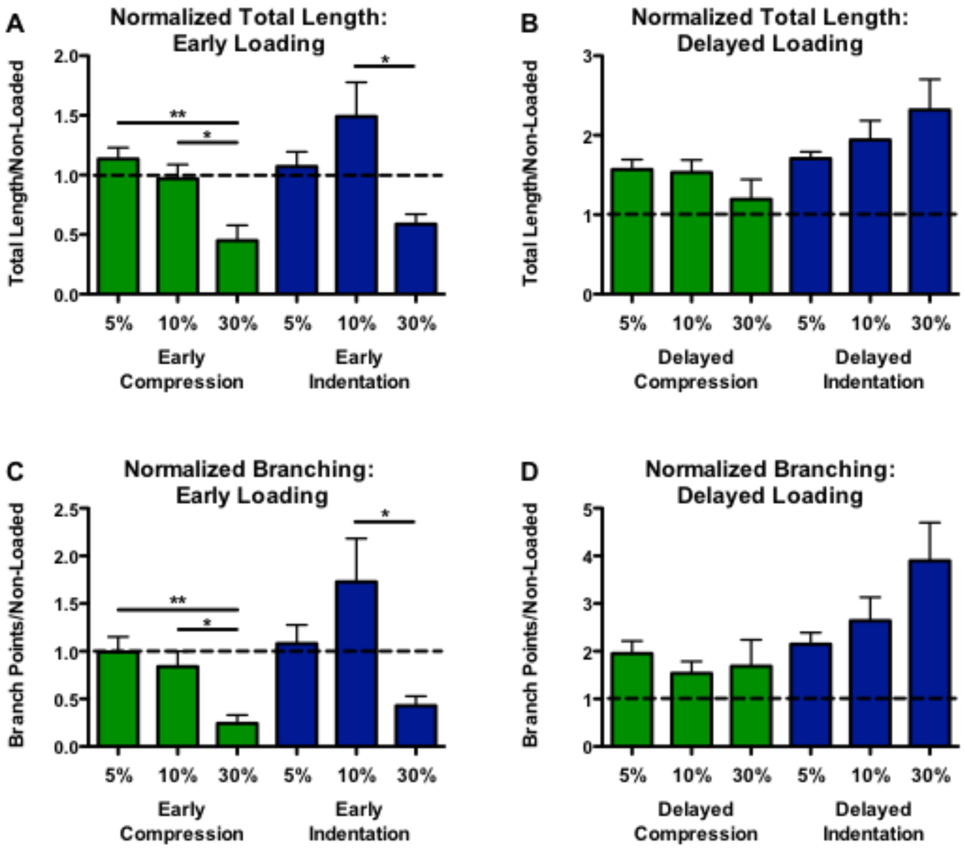


**Figure S2.** Vessel network length and branching under 5%, 10%, and 30% strain normalized to non-loaded group. 1-way ANOVA, * $p<0.05$, ** $p<0.01$. n=6/group.

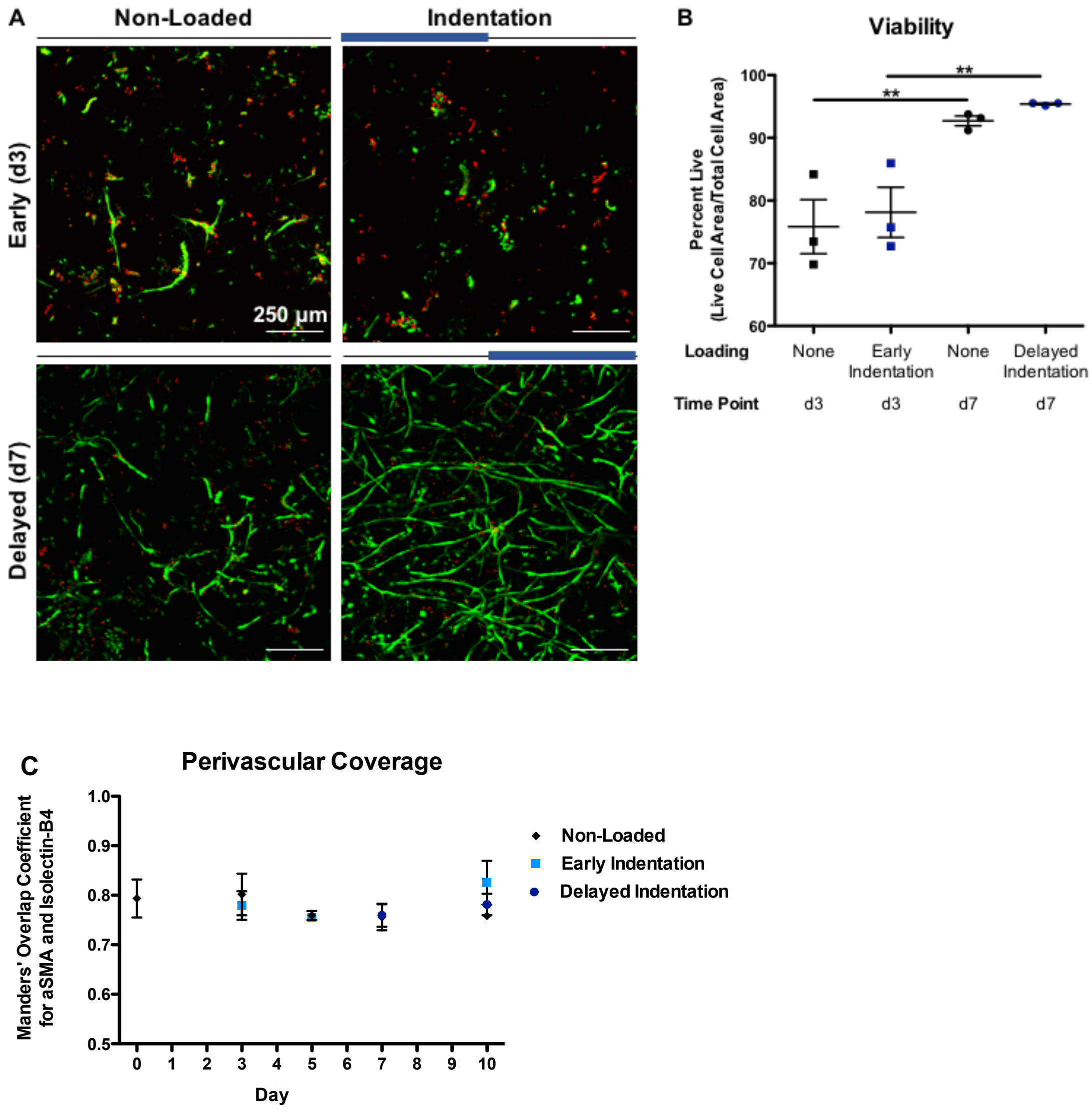


**Figure S3.** Cell viability under early and delayed 30% indentation. A) Representative maximum intensity z-projections (25 µm depth) of calcein (green; live) and ethidium homodimer (red; dead) stained microvascular fragments at days 3 (early) and 7 (delayed). B) Image-based quantification of live/dead stain. 2-way ANOVA, ** Bonferroni post hoc p<0.01. No effect of loading. No interaction effect. n=3 random images/gel, n=3 gels/group/time point. C) Image-based quantification of perivascular coverage (pictured in Figure 3 B). 2-way ANOVA, no significant differences due to loading or time. n=3 gels/group/time point.

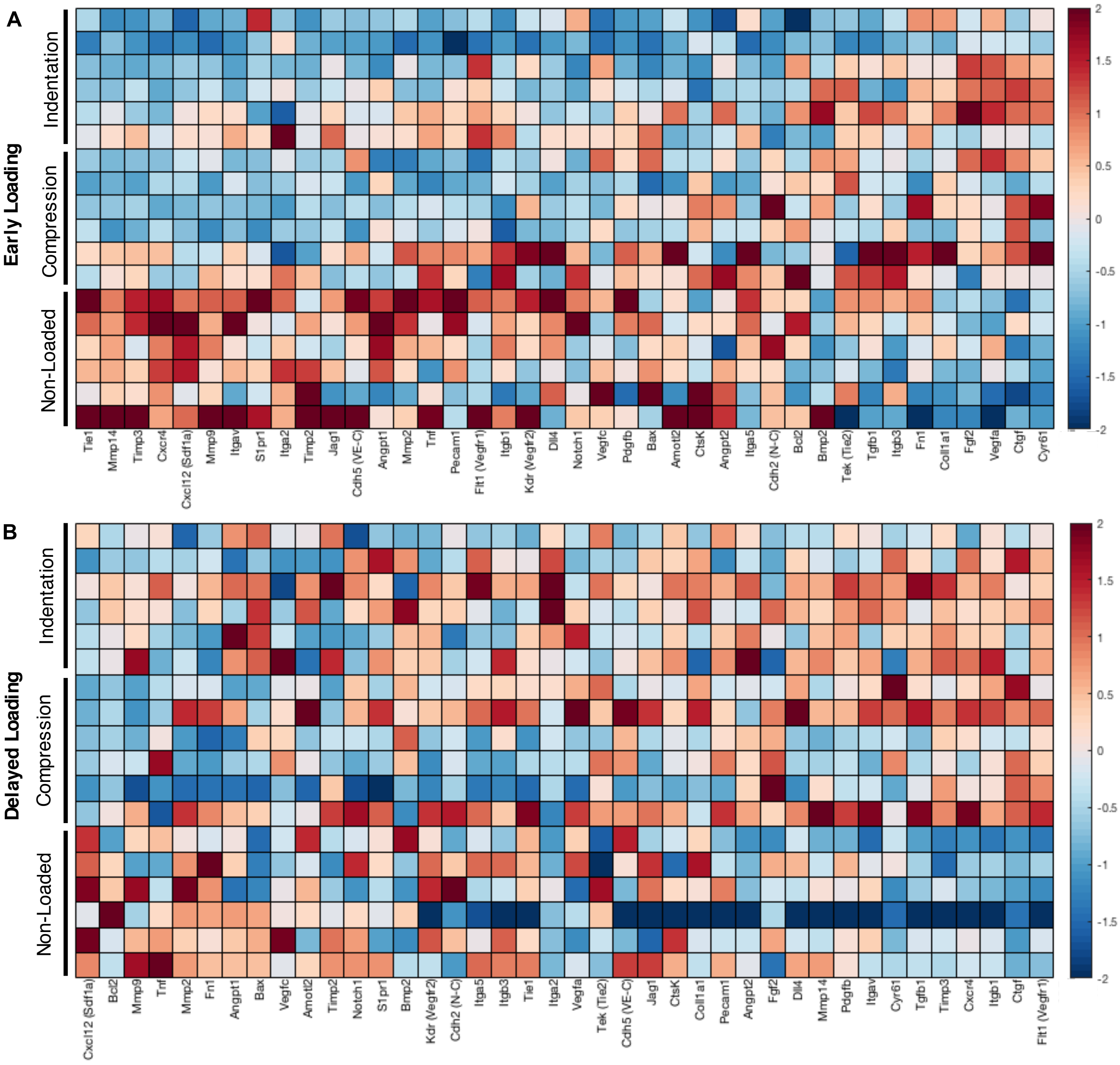


**Figure S4.** Z-scored gene expression data under A) early and B) delayed dynamic 30% strain at 1 Hz.

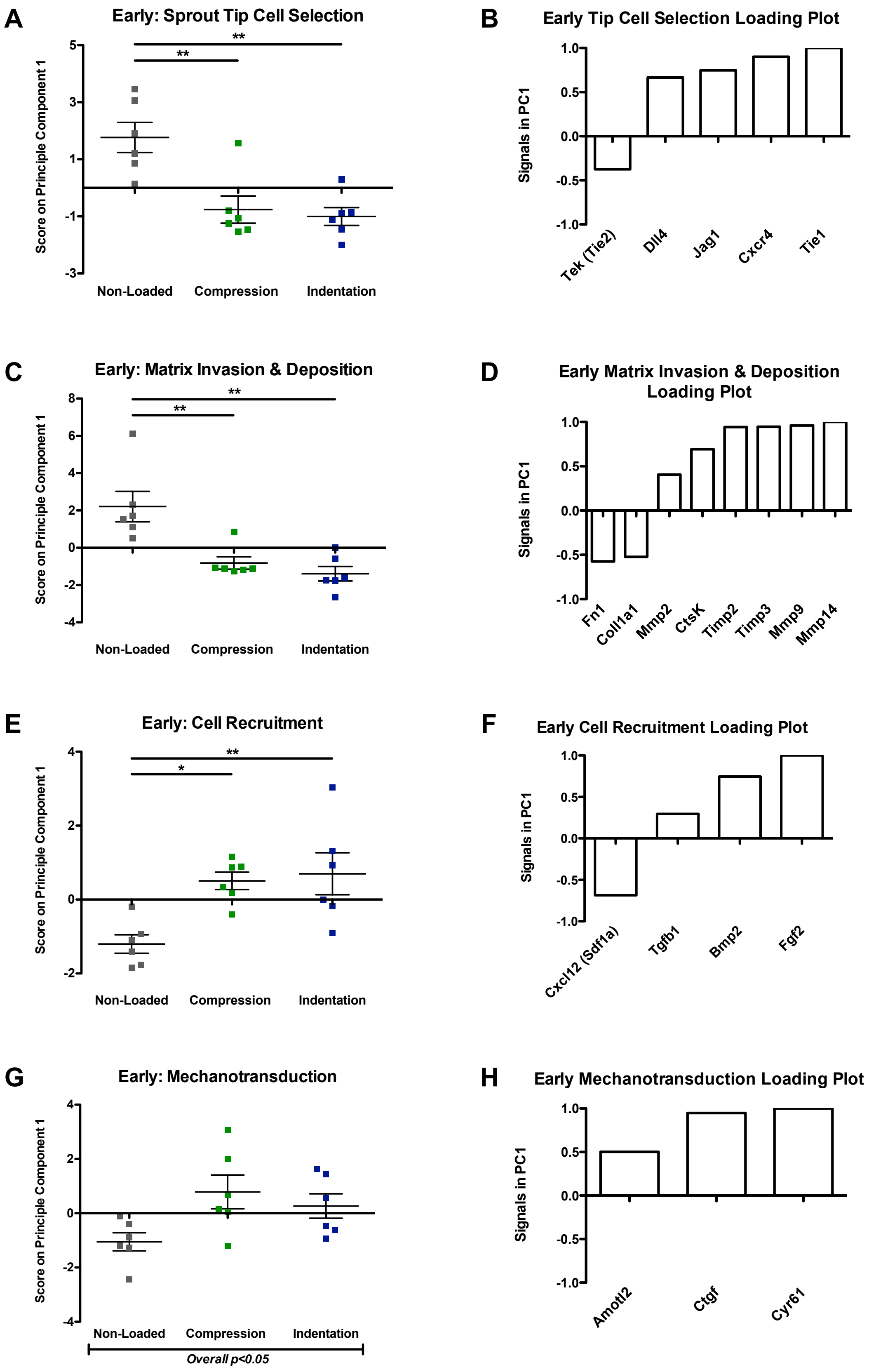
A
Early: Sprout Tip Cell Selection
Score on Principle Component 1
Non-Loaded
Compression
Indentation
B
Early Tip Cell Selection Loading Plot
Signals in PC1
Tek (Tie2)
Dll4
Jag1
Cxcr4
Tie1
C
Early: Matrix Invasion & Deposition
Score on Principle Component 1
Non-Loaded
Compression
Indentation
D
Early Matrix Invasion & Deposition Loading Plot
Signals in PC1
Fn1
Coll1a1
Mmp2
CtsK
Timp2
Timp3
Mmp9
Mmp14
E
Early: Cell Recruitment
Score on Principle Component 1
Non-Loaded
Compression
Indentation
F
Early Cell Recruitment Loading Plot
Signals in PC1
Cxcl12 (Sdf1a)
Tgfb1
Bmp2
Fgf2
G
Early: Mechanotransduction
Score on Principle Component 1
Non-Loaded
Compression
Indentation
Overall p<0.05
H
Early Mechanotransduction Loading Plot
Signals in PC1
Amotl2
Ctgf
Cyr61

**Figure S5.** Gene Sets significantly affected by early 30% strain at 1 Hz as determined by PCA. Score on principal component 1 and corresponding loading plots of gene sets known to be involved in sprout tip cell selection (A-B), matrix invasion and deposition (C-D), cell recruitment (E-F), and mechanotransduction (G-H). 1-way ANOVA, $*p<0.05$, $**p<0.01$.

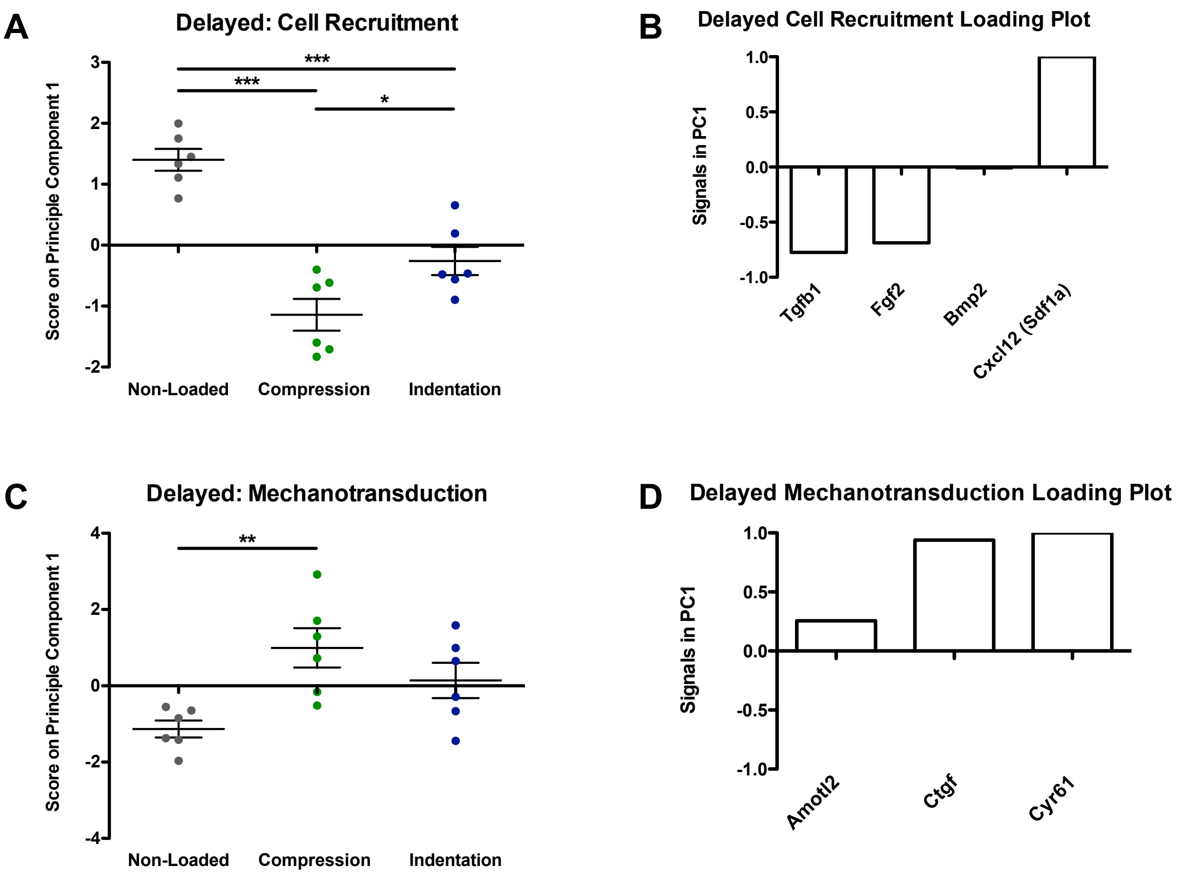


**Figure S6.** Gene Sets significantly affected by delayed 30% strain at 1 Hz as determined by PCA. Score on principal component 1 and corresponding loading plots of gene sets known to be involved in cell recruitment (A-B), and mechanotransduction (C-D). 1-way ANOVA, *p<0.05, **p<0.01,***p<0.001.

### Downregulated by Early Loading

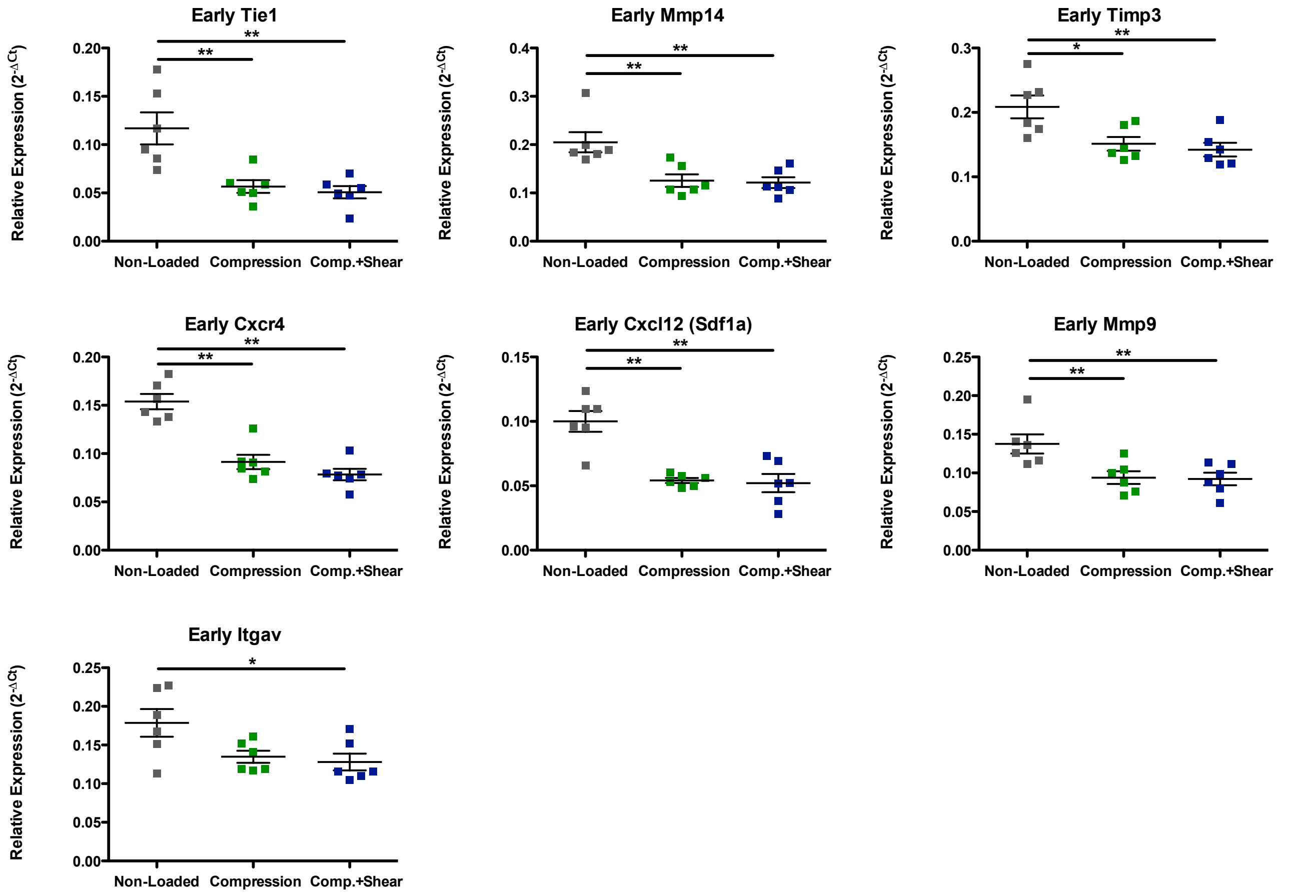


### Upregulated by Early Loading

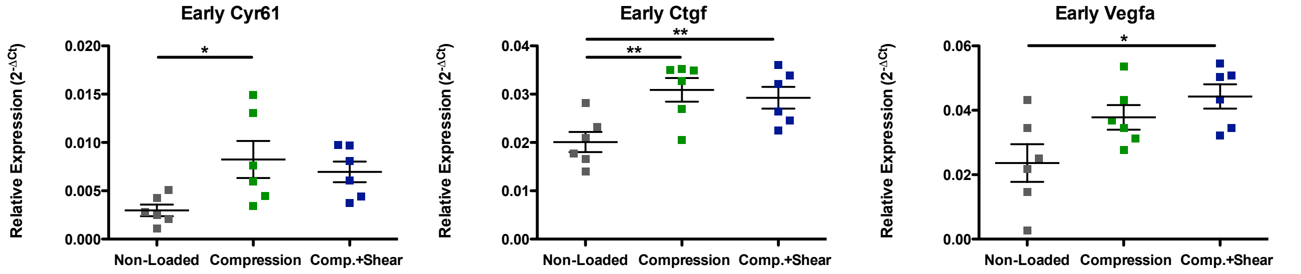


**Figure S7.** Individual genes significantly affected by early 30% strain at 1 Hz.

1-way ANOVA, Bonferroni post hoc $*p<0.05$, $**p<0.01$. n=6/group.

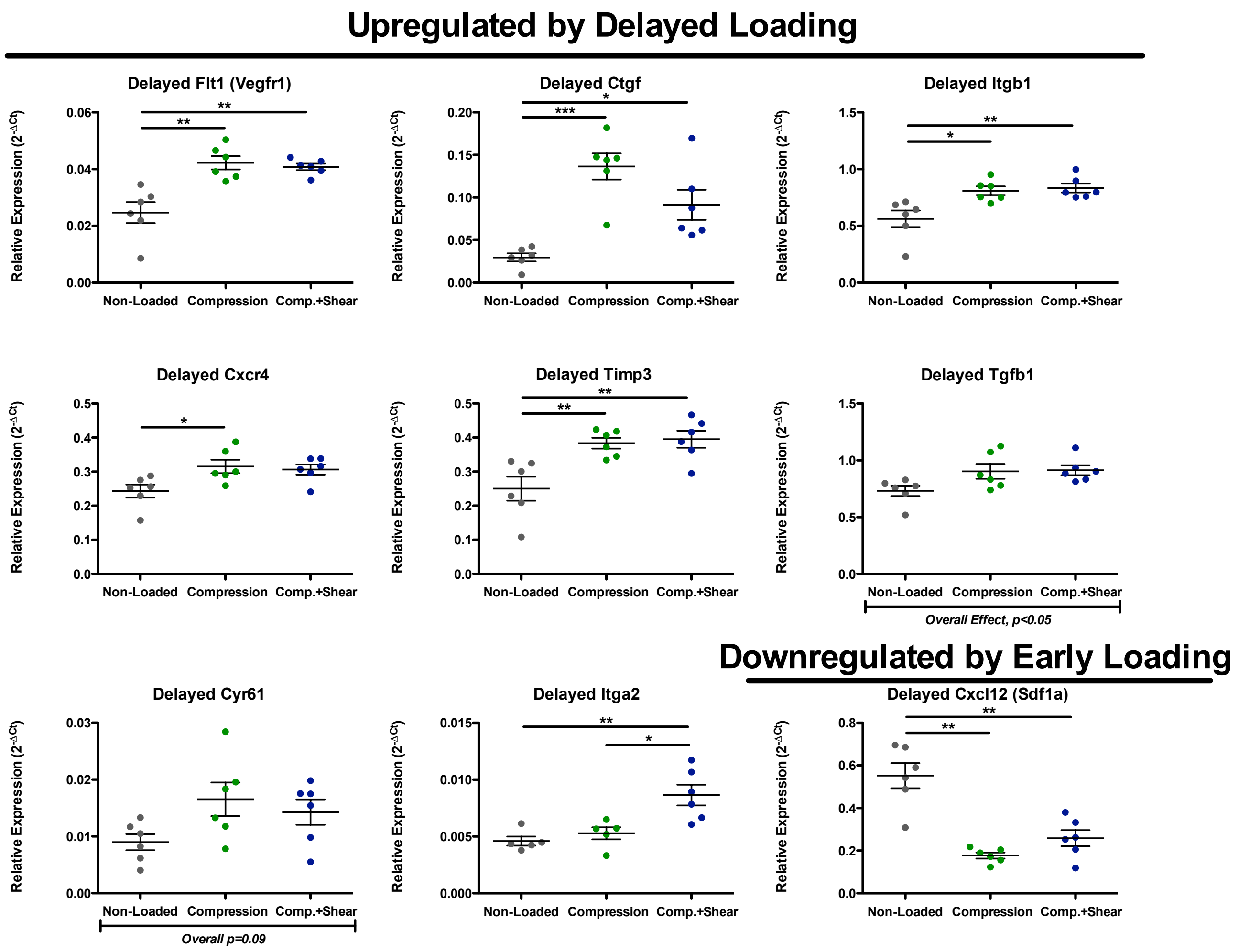


**Figure S8.** Individual genes significantly affected by delayed 30% strain at 1 Hz. 1-way ANOVA, Bonferroni post hoc $*p<0.05$, $**p<0.01$, $***p<0.001$. n=6/group.

**Supplemental Table 1.** List of Genes Measured with Taqman Probes Using Fluidigm System.

| Stage of Angiogenesis | Gene | Protein Encoded | Taqman Reference Number |
|---|---|---|---|
| Sprout Tip Cell Selection | *Notch1* | Notch 1 | Rn01758633_m1 |
| Sprout Tip Cell Selection | *Jag1* | Jagged 1 | Rn00569647_m1 |
| Sprout Tip Cell Selection | *Dll4* | Delta-like ligand 4 | Rn01512886_m1 |
| Sprout Tip Cell Selection | *Tie1* | Tie 1 | Rn01417182_m1 |
| Sprout Tip Cell Selection | *Tek (Tie2)* | Tie 2 | Rn01433346_m1 |
| Sprout Tip Cell Selection | *Cxcr4* | C-X-C Motif Chemokine Receptor 4 | Rn01483207_m1 |
| Vessel (De)stabilization and Growth | *Vegfa* | Vascular endothelial growth factor a | Rn01511602_m1 |
| Vessel (De)stabilization and Growth | *Vegfc* | Vascular endothelial growth factor c | Rn01488076_m1 |
| Vessel (De)stabilization and Growth | *Flt1 (Vegfr1)* | Vascular endothelial growth factor receptor 1 | Rn01409533_m1 |
| Vessel (De)stabilization and Growth | *Kdr (Vegfr2)* | Vascular endothelial growth factor receptor 2 | Rn00564986_m1 |
| Vessel (De)stabilization and Growth | *Angpt1* | Angiopoietin 1 | Rn01504818_m1 |
| Vessel (De)stabilization and Growth | *Angpt2* | Angiopoietin 2 | Rn01756774_m1 |
| Vessel (De)stabilization and Growth | *Pdgfb* | Platelet derived growth factor subunit b | Rn01502596_m1 |
| Matrix Invasion and Deposition | *Mmp2* | Matrix metalloproteinase 2 | Rn01538170_m1 |
| Matrix Invasion and Deposition | *Mmp9* | Matrix metalloproteinase 9 | Rn00579162_m1 |
| Matrix Invasion and Deposition | *Mmp14* | Matrix metalloproteinase 14 | Rn00579172_m1 |
| Matrix Invasion and Deposition | *Timp2* | Tissue inhibitor of metalloproteinase 2 | Rn00573232_m1 |
| Matrix Invasion and Deposition | *Timp3* | Tissue inhibitor of metalloproteinase 3 | Rn00441826_m1 |
| Matrix Invasion and Deposition | *CtsK* | Cathepsin K | Rn00580723_m1 |
| Cell Adhesion and Migration | *Cdh5* | Vascular endothelial cadherin | Rn01536708_m1 |
| Cell Adhesion and Migration | *Cdh2* | Neural cadherin | Rn00580099_m1 |
| Cell Recruitment | *Bmp2* | Bone morphogenetic protein 2 | Rn00567818_m1 |
| Cell Recruitment | *Cxcl12 (Sdf1a)* | Stromal derived factor 1 alpha | Rn00573260_m1 |
| Cell Recruitment | *Fgf* | Fibroblast growth factor | Rn00570809_m1 |

| | | | |
|---|---|---|---|
| Cell Recruitment | *S1pr1* | Sphingosine 1-phosphate receptor 1 | Rn02758712_s1 |
| Cell Recruitment | *Tgfb1* | Transforming growth factor b | Rn00572010_m1 |
| Cell Adhesion and Migration | *Pecam1* | Platelet endothelial cell adhesion molecule 1 | Rn01467262_m1 |
| Cell Adhesion and Migration | *Itga5* | Integrin alpha 5 | Rn01761831_m1 |
| Cell Adhesion and Migration | *Itgb1* | Integrin beta 1 | Rn00566727_m1 |
| Cell Adhesion and Migration | *Itgav* | Integrin alpha v | Rn01485633_m1 |
| Cell Adhesion and Migration | *Itgb3* | Integrin beta 3 | Rn00596601_m1 |
| Cell Adhesion and Migration | *Itga2* | Integrin alpha 2 | Rn01489315_m1 |
| Inflammation and Apoptosis | *Il12* * | Interleukin 12 | Rn00584538_m1 |
| Inflammation and Apoptosis | *Il10* * | Interleukin 10 | Rn01483988_g1 |
| Inflammation and Apoptosis | *Tnfa* | Tumor necrosis factor alpha | Rn01525859_g1 |
| Matrix Invasion and Deposition | *Fn* | Fibronectin | Rn00569575_m1 |
| Matrix Invasion and Deposition | *Coll1a1* | Collagen type 1 alpha 1 | Rn01463848_m1 |
| Mechanotransduction | *Amotl2* | Angiomotin-like 2 | Rn01446301_m1 |
| Mechanotransduction | *Ankrd1* * | Ankyrin repeat domain 1 | Rn00566329_m1 |
| Mechanotransduction | *Ctgf* | Connective tissue growth factor | Rn01537279_g1 |
| Mechanotransduction | *Cyr61* | Cyr61 | Rn00580055_m1 |
| Inflammation and Apoptosis | *Bax* | Bcl-2-associated X protein | Rn01480161_g1 |
| Inflammation and Apoptosis | *Bcl2* | B-cell lymphoma 2 | Rn99999125_m1 |
| Housekeeping | *Gapdh* | Glyceraldehyde 3-phosphate dehydrogenase | Rn01775763_g1 |
| Housekeeping | *Bactin* | Beta-actin | Rn00667869_m1 |
| Housekeeping | *Hprt1* | Hypoxanthine-guanine phosphoribosyltransferase | Rn01527840_m1 |
| Housekeeping | *Ubc* | Poly-ubiquitin C | Rn01499642_m1 |
| Housekeeping | *Ppia* | Peptidylprolyl isomerase A | Rn00690933_m1 |
| ** Indicates gene expression was below the limit of detection and thus excluded from analyses.* | | | |
